# Structural basis for coupling of the WASH subunit FAM21 with the endosomal SNX27-Retromer complex

**DOI:** 10.1101/2023.08.15.553351

**Authors:** Qian Guo, Kai-en Chen, Manuel Gimenez-Andres, Adam P. Jellett, Ya Gao, Boris Simonetti, Meihan Liu, Chris M. Danson, Kate J. Heesom, Peter J. Cullen, Brett M. Collins

## Abstract

Endosomal membrane trafficking is mediated by specific protein coats and formation of actin-rich membrane domains. The Retromer complex coordinates with sorting nexin (SNX) cargo adaptors including SNX27, and the SNX27–Retromer assembly interacts with the WASH complex which nucleates actin filaments establishing the endosomal reycling domain. Crystal structures, modelling, biochemical and cellular validation reveal how the FAM21 subunit of WASH interacts with both Retromer and SNX27. FAM21 binds the FERM domain of SNX27 using acidic-Asp-Leu-Phe (aDLF) motifs similar to those found in the SNX1 and SNX2 subunits of the ESCPE-1 complex. Overlapping FAM21 repeats and a specific Pro-Leu containing motif bind three distinct sites on Retromer involving both the VPS35 and VPS29 subunits. Mutation of the major VPS35-binding site does not prevent cargo recycling, however it partially reduces endosomal WASH association indicating that a network of redundant interactions promote endosomal activity of the WASH complex. These studies establish the molecular basis for how SNX27–Retromer is coupled to the WASH complex via overlapping and multiplexed motif-based interactions required for the dynamic assembly of endosomal membrane recycling domains.

**Significance Statement:** Cell surface transmembrane proteins are regulated by a constant cycle of internalization and recycling from intracellular endosomal compartments. The Retromer protein complex and the sorting nexin adaptor protein play a critical role in the retrieval of hundreds of proteins responsible for ion transport, glucose metabolism, neurotransmission, and other cell functions. We have defined the mechanism by which both Retromer and SNX27 engage the actin-nucleating complex called WASH through multiple repeated sequences in the subunit FAM21. Dysfunction in WASH, Retromer and SNX27 are implicated in various disorders including Parkinson’s disease, Alzheimer’s disease, hereditary spastic paraplegia, and this work provides important insights into the assembly of these essential endosomal sorting machineries.

## INTRODUCTION

Maintenance of appropriate levels of endosomal sorting of transmembrane ‘cargo’ proteins is essential for cellular function (1). Mammalian Retromer is an evolutionarily conserved protein complex composed of VPS35, VPS29, and VPS26 (two isoforms VPS26A and VPS26B) and is a central regulator of endosomal sorting across eukaryotes (2, 3). More specifically, it functions together with diverse accessory proteins and cargo adaptors to coordinate endosomal recruitment and sequence-specific cargo recognition with the assembly of endosomal sub-domains for the biogenesis of tubulovesicular transport carriers. Sorting nexin 27 (SNX27) is a Retromer cargo adaptor that binds to specific PSD95-DLG-ZO1 (PDZ) domain binding motif (PDZbm) containing cargo to promote their Retromer-dependent endosome-to-cell surface recycling (4–8).

Both Retromer and SNX27 associate with the Wiskott-Aldrich Syndrome Protein and SCAR Homolog (WASH) complex, which by driving localised actin polymerisation plays an essential role in endosomal sorting and transport (7–13). The WASH complex is a pentamer of WASHC1, FAM21 (WASHC2), WASHC3 (coiled-coil domain-containing protein 53 (CCDC53)), WASHC4 (KIAA1033; WASH-interacting protein; SWIP) and WASHC5 (Strumpellin; KIAA0196) (9, 10, 14, 15). The WASHC1 subunit is a member of the Wiskott-Aldrich Syndrome Protein (WASP) family (14, 16), and the presence of a C-terminal *<u>v</u>*erprolin homology sequence, *<u>c</u>*onnecting sequence, and *<u>a</u>*cidic sequence (VCA) domain allows WASHC1 to activate the actin-related protein 2/3 (ARP2/3) complex to stimulate formation of branched F-actin networks (17). The ARP2/3 complex plays a number of roles in regulating endocytosis and subsequent endosomal sorting and transport processes (18–20).

During endosomal cargo sorting and transport, the WASH complex is essential for the fission of tubules during the biogenesis of transport carriers. Endosomal tubulation is exaggerated in cells lacking the WASH complex (10) and WASH depletion leads to the observation of long endosomal tubules aligned along microtubules (9). It is thought that WASH promotes the scission of cargo-containing tubules via generating F-actin-driven force at the endosomal membrane. Mutations in WASH complex subunits can result in neurological pathologies, such as hereditary spastic paraplegia, inherited intellectual disability, and late-onset Alzheimer’s disease (15, 21–24).

Retromer regulates the recruitment of a population of the WASH complex to endosomal membranes (12, 13), while binding of WASHC4 to membrane phosphoinositide lipids can promote Retromer-independent endosomal association. Native immunoprecipitations provided the initital evidence for the WASH complex interacting with Retromer, while yeast two-hybrid assays suggested direct associations between VPS35, FAM21 and WASHC1 (12). Suppression and knock-out of Retromer partly reduces WASH endosomal association (12, 25), while other members of the endosomal protein sorting system are unaffected, such as SNX1 and the RAB5 effector EEA1 (12). Structurally, FAM21 has a 200 amino acid “head” domain that interacts with the WASHC5 and WASHC1 proteins and is crucial for the formation and stabilisation of the WASH complex (13, 14). Following the head domain is a large (around 1100 amino acids) unstructured “tail” domain that contains 21 repeating motifs composed of Leu-Phe and several acidic residues (Asp or Glu) dubbed LFa motifs (26). The enormous unstructured tail of FAM21 is important for membrane association (10), and it was shown that the LFa motifs can bind to the VPS35 subunit of Retromer with the very C-terminal motifs being most crucial for the interaction (26, 27). Overexpression of the FAM21 tail can displace the WASH complex on the endosome by competing for binding to VPS35 and potentially other effectors (13, 26, 27). Although the VPS35 subunit can associate with the FAM21 tail directly in yeast two-hybrid experiments and in isothermal titration calorimetry (ITC) assays (13, 26), binding is enhanced by other subunits when VPS35 is assembled into the functionally active Retromer complex (27). The interaction of Retromer with WASH via FAM21 is thought to link branched actin polymerisation to cargo protein sorting and trafficking activity, and the long unstructured nature of the FAM21 tail combined with the many LFa repeats likely promotes both avidity for Retromer and flexibility in the relative orientations for dynamic endosomal membrane sorting.

The SNX27 Retromer cargo adaptor was also reported to associate with the WASH complex by coimmunoprecipitation (7). Responsible for recycling over 400 specific transmembrane cargoes from endosomes to the plasma membrane (5, 7, 28–30), SNX27 is a peripheral membrane protein containing, from N-termini to C-termini, PDZ, PX and FERM domains that mediate cargo selection, Retromer recruitment and protein-lipid interactions (4, 30, 31). Notably, SNX27 can bind via its FERM domain to acidic DLF (aDLF) motifs in the N-terminal disordered region of SNX1 and its homologue SNX2 within the BAR domain-containing ESCPE-1 complex (32, 33).

Here we show that the FAM21 subunit of WASH coordinates the interactions with both Retromer and SNX27 via very different mechanisms but incorporating distinct overlapping elements of the repeat sequences in its extended C-terminal domain. Crystal structures, modelling, biochemical and cellular experiments show that the multiple redundant acidic-Asp-Leu-Phe (aDLF) repeat sequences in FAM21 bind to a conserved basic pocket in the SNX27 FERM domains and are essentially identical to the aDLF sequences found in the endosomal proteins SNX1 and SNX2 (32). Overlapping repeated motifs in FAM21 with the consensus sequence [L/I]F<ED<_3-9_[L/I]F bind two conserved sites towards the C-terminus of Retromer subunit VPS35. These lie opposite to the VSP29 binding site and adjacent to the dimerising surface required for assembling Retromer arches on the membrane. In addition, the last repeat of FAM21 contains a specific Asp-Pro-Leu sequence that binds a third site on VPS29 that is the same region bound by other interactors such as TBC1D5 and ANKRD27. Mutations in these regions perturb FAM21 interaction *in vitro* and in cells, however single mutations in VPS35 are not sufficient to block Retromer-mediated trafficking, consistent with a model whereby the WASH complex engages the endosome and associated proteins through multiple related but redundant motif-mediated interactions.

## RESULTS

### The FAM21 tail binds to both VPS35 and VPS29 through its repeat sequences

A primary characteristic of FAM21 is the long disordered tail enriched in acidic residues and containing 21 copies of a repeated [I/L]F sequence with C-terminal acidic residues (**Figures 1A and 1B**). This was previously termed the LFa motif and shown to bind directly to the Retromer subunit VPS35 with the final three repeats, 19-21, showing the strongest binding affinity (26). However, suggesting that VPS29 also plays a role in binding FAM21, we recently noted that binding of Retromer and GST-tagged FAM21 composed of repeats 19, 20 and 21 (GST-FAM21_R19-R21_) was slightly reduced in the presence of a macrocyclic peptide that binds via a PL-containing sequence to a conserved site on the VPS29 subunit (34). To examine this, we first tested the binding between Retromer and peptides from human FAM21 repeat 19 (FAM21_R19_), repeat 20 (FAM21_R20_) and repeat 21 (FAM21_R21_). Using ITC, we found FAM21_R21_ alone was able to bind to either Retromer, to the subcomplex composed of VPS29 and VPS35 residues 481 to 796 (VPS29–VPS35_481-796_), as well as VPS29 alone (**Figure 1C; Table S1**). In contrast, FAM21_R19_ and FAM21_R20_ peptides showed a relatively weak Retromer interaction and no interaction at all with VPS29 (**Figure 1C)**. A stronger Retromer interaction was observed when we tested the GST-tagged construct combining repeats 19 and 20 (GST-FAM21_R19-R20_), suggesting some additive effect of these two sequences (**Figure 1D)**. Notably the FAM21_R21_ interaction with VPS29 on its own could be blocked by the addition of inhibitory cyclic peptide RT-D1 (**Figure 1E**) (34), indicating that FAM21_R21_ likely binds to the same conserved VPS29 surface. Unlike other repeats FAM21 repeat 21 also contains a conserved DPL sequence next to the [I/L]Fa motif (**Figure 1B; Figure S1**) and it is this sequence that binds to VPS29 while other repeats bind to VPS35.

**Figure 1.**
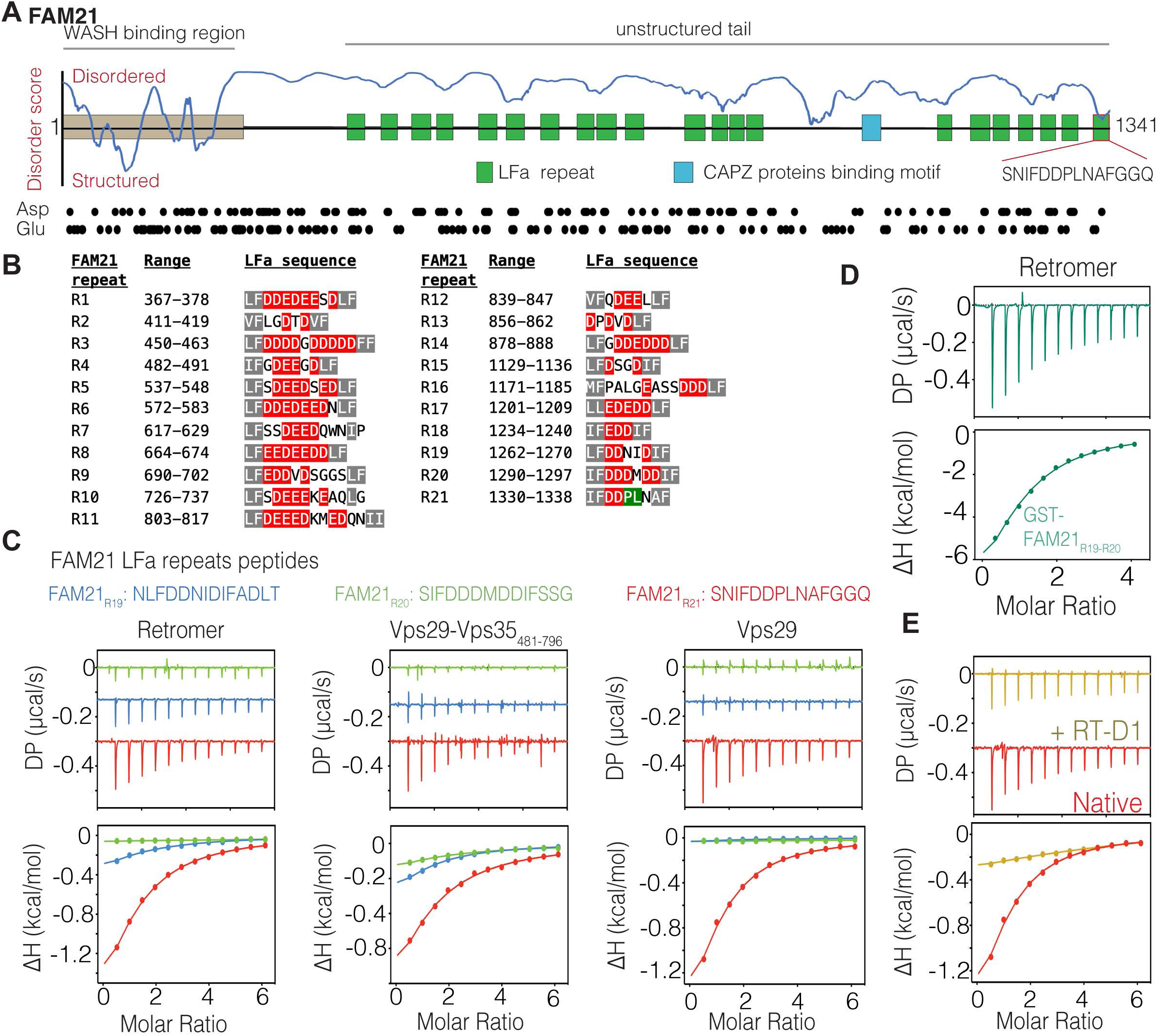
Retromer VPS29 is responsible for binding to repeat 21 of FAM21. (**A**) Schematic diagram showing features of human FAM21, and highlighting the 21 LFa repeats in the unstructured tail. (**B**) Sequences of the 21 LFa repeats within the FAM21 tail. Conserved [I/L]F residues are shown in grey and acidic residues are in red. The PL residues of repeat 21 are highlighted in green. (**C**) ITC thermograms showing the titration of Retromer, the VPS29–VPS35_481-796_ subcomplex, and VPS29 alone with FAM21_R19_ (blue); FAM21_R20_ (green); and FAM21_R21_ (red) peptides. (**D**) Retromer shows binding to GST-FAM21_R19-R20_. (**E**) VPS29 and FAM21_R21_ peptide interaction can be blocked by the addition of the PL-sequence containing cyclic peptide RT-D1. All ITC graphs show the integrated (bottom) and normalized raw data (top) fit with a single-site binding model.

### Structural basis for the interaction of FAM21 LFa repeats with VPS35

Our data suggests that FAM21 LFa repeats 19 and 20 interact with VPS35 while the divergent DPL-containing repeat 21 can bind VPS29. To understand the molecular basis of how the LFa repeating units bind to Retromer, we attempted to co-crystallize the VPS29–VPS35_481-796_ subcomplex with FAM21_R19_ and FAM21_R20_ peptides. The structure of VPS29–VPS35_481-796_ in complex with FAM21_R20_ peptide was successfully solved at 3 Å resolution with two VPS29–VPS35_481-796_ molecules in the asymmetric unit (**Figure 2A; Table S2**). Although the structure was solved at relatively low resolution, we could observe additional electron density on the surface of VPS35 representing the FAM21_R20_ peptide (**Figure 2B**). The electron density for the FAM21_R20_ peptide is located on a conserved positively charged groove of VPS35 between helices 23 and 25 surrounded by K552, K555, K556 and K559 (**Figures 2B and 2C**). To aid in assigning the FAM21_R20_ residues to the structure as well as investigate the potential binding surface of repeat 19, we used AlphaFold2 modelling to predict complexes of different variations of FAM21 sequences bound to Retromer. These predictions consistently identified two separate binding sites on VPS35 for the FAM21_R19-R20_ LFa repeats (**Figures 2D and 2E**). Inspecting these models closely, we found that repeats 19 and 20 are predicted to bind two distinct positively charged grooves at the C-terminal end of the VPS35 surface that we refer to as Site 1 and Site 2 respectively (**Figure 2E**). The X-ray crystallographic difference density for the FAM21_R20_ peptide aligns with Site 2 and matches well to the residues ^1296^DDIF^1299^ of repeat 20 predicted in the model (**Figure 2F**). In particular I1298 and F1299 are positioned to interact with K555, K556 and K559 of VPS35 (**Figure 2F**). A notable feature of the predicted structure is that in addtion to the ^1296^DDIF^1299^ residues the repeated element engages VPS35 via an additional IF sequence interspersed with acidic residues ^1290^<u>IF</u>DDDMDD<u>IF</u>^1299^ (**Figure 2F**). Similarly the predictions indicate that the second R19 sequence can bind to Site 1 with a similar dual [I/L]F motif with interspersed acidic residues ^1262^<u>LF</u>DDNID<u>IF</u>^1270^ recognising a second positively charged groove composed of K701, K705 and K749 (**Figure 2E**). With the combination of predictions and experimental structure we propose that VPS35 possesses two distinct binding sites for low affinity binding of the multiple LFa sequences in FAM21 (**Figure 1B**)(26). These structures indicate the the LFa motif ideally possesses two [I/L]F sequences with an interveining stretch of Asp and Glu side chains [I/L]F[E/D]_n_[I/L]F.

**Figure 2.**
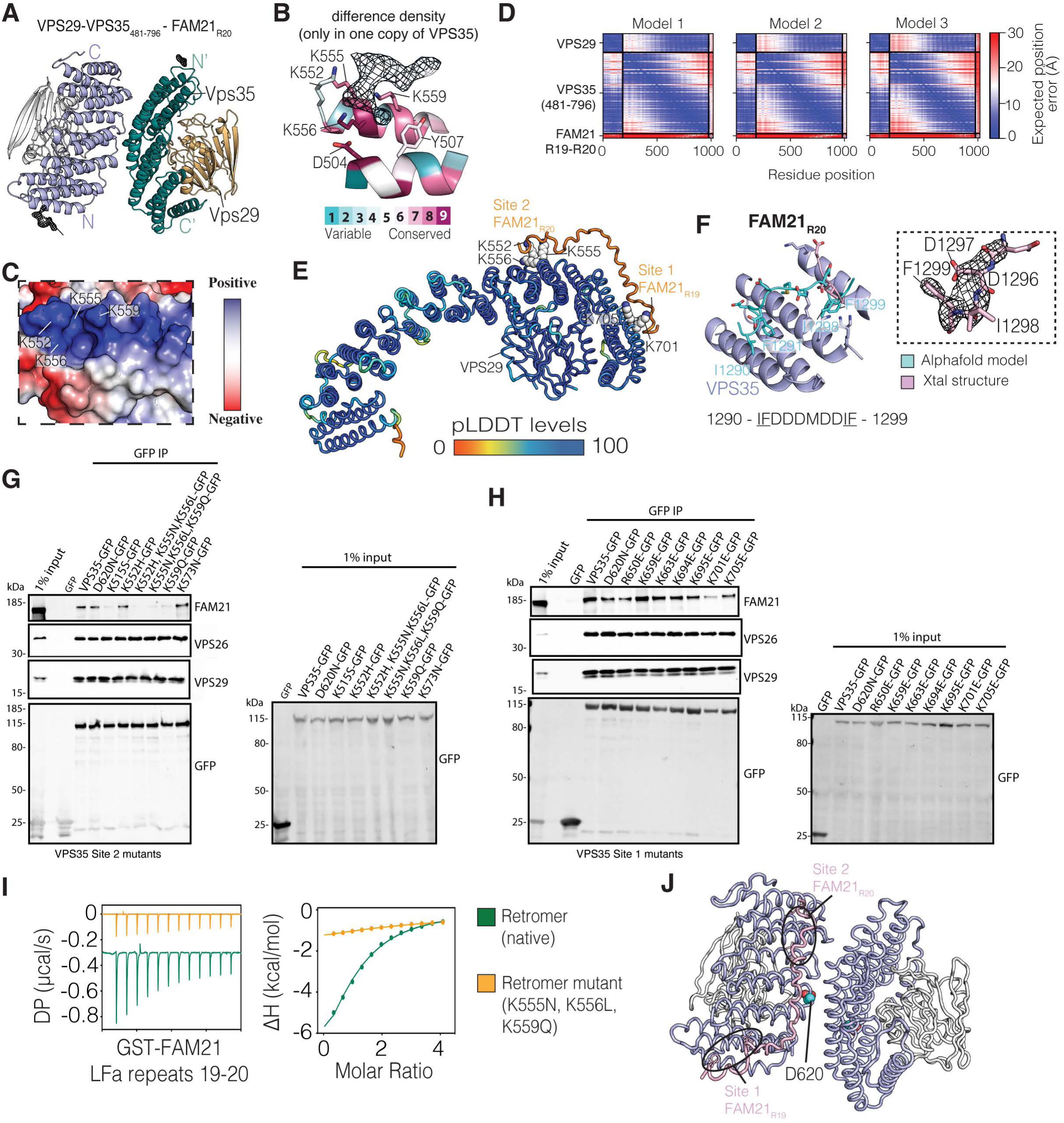
Structural basis for the interaction of LFa repeats with Retromer subunit VPS35. (**A**) Overall crystal structure of VPS29–VPS35_481-796_ in complex with FAM21_R20_ peptides showing both complexes from the asymmetric unit. (**B**) VPS35 is highly conserved in the FAM21_R20_ binding region. The density corresponding to the FAM21_R20_ peptide is shown in mesh using 2Fo–Fc map contoured at 1σ. Sequence conservation of VPS35 was mapped with CONSURF (75). (**C**) The electrostatic surface map showing FAM21_R20_ binding location on VPS35. (**D**) Predicted alignment Error (PAE) plots of VPS29–VPS35_481-796_– FAM21_R19-R20_ complexes generated by Alphafold2. (**E**) The predicted structure highlighting the per-residue confidence score (pLDDT) between 0 to 100. Regions showing low pLDDT score (green and orange) are likely to be unstructured. (**F**) Alignment of the Alphafold2 model of FAM21_R20_ (cyan) and the crystal structure (VPS35 in light purple, FAM21_R20_ as light pink) highlighting the key interacting residues between FAM21_R20_ and VPS35. The insert shows the refined electron density of the FAM21_R20_ peptide observed in the crystal structure. (**G**) Co-immunoprecipitation (co-IP) of C-terminal GFP-tagged VPS35mutants in the Site 2 binding pocket in HEK293T cells. (**H**) Co-IP of C-terminal GFP-tagged VPS35 and mutants in the Site 1 binding pocket. Quantitation of these Western blots is shown in **Figure S2.** (**I**) ITC analysis showing the binding of GST-FAM21_R19-R20_ with either native Retromer or VPS35 (K555N/K556L/K559Q) Site 2 mutant. All ITC graphs show the integrated (bottom) and normalized raw data (top) fit with a single-site binding model. (**J**) The locations of the Site 1 and Site 2 FAM21 repeat binding sites are shown in the context of the VPS35-mediated Retromer dimer assembly derived from cryoelectron tomography (37). The location of the D620N mutation linked to Parkinson’s Disease is also indicated in cyan spheres.

To further validate if both positively charged grooves of VPS35 are responsible for binding to FAM21 we performed co-immunoprecipitation with structurally targeted site-directed mutations of VPS35. Importantly, all mutants retained the ability to assemble into the Retromer heterotrimeric complex **(Figure 2G and 2H; Figure S2)**. Consistent with our structural data, binding to FAM21 was significantly affected by the multiple mutations around the repeat 20 binding groove including K552, K555, K556 and K559 (**Figure 2G; Figure S2**). The strongest defect in FAM21 binding was observed with the triple mutant composed of K555, K556 and K559 (**Figures 2G and 2I**; **Figure 2G; Figure S2**), the residues known to form contact with the LFa motif in Site 2. In addition, mutations of other positively charged residues surrounding the core [I/L]F dipeptide including K515S also showed significant defects in binding. It is likely that these residues together form a large positively charged patch on VPS35 for the initial recognition of the acidic repeats of FAM21. A similar defect was also observed in the K701E mutant on the second positively charged groove of Site 1 (**Figure 2H**). Notably, we also observed a slight defect in FAM21 binding when we used the VPS35 D620N mutant, the mutation known to be involved in the Parkinson diseases (35, 36). The mutation is distal from the FAM21_R19_ and FAM21_R20_ binding sites but lies next to the VPS35-mediated Retromer dimerization interface as seen in the Retromer-SNX3 and Retromer-SNX-BAR tubular coat complexes (37) (**Figure 2J**). It is possible that the slight defect in FAM21 binding is indirectly caused by interference with Retromer dimerization.

### Repeat 21 binds to VPS29 through a classic PL motif interaction

To better understand the association of FAM21 repeat 21 with Retromer we next crystallised the VPS29– VPS35_481-796_ complex with the FAM21_R21_ repeat and determined its crystal structure at a modest 3.4 Å resolution. The structure reveals two VPS29–VPS35_481-796_ molecules in the asymmetric unit. Notably, the dimerization interface in this structure differ from that observed in the FAM21_R20_ bound structure (**Figure 3A**). Despite the low resolution, we could still observe additional electron density around the conserved hydrophobic pocket of VPS29 (**Figure 3A**). To unambiguously assign the key residues involve in the interaction, we then successfully solved the structure of the VPS29–FAM21_R21_ peptide complex at 2.0 Å resolution (**Figure 3B**). The improved density allowed us to build the PL motif and upstream residues. According to the structure, we could unambiguously observe the density up to D1332, suggesting that the signature [I/L]Fa dipeptide located at position 1330 and 1331 was not involved in the binding (**Figure 3C**). In direct contrast to the FAM21_R20_ structure bound with VPS35, the FAM21_R21_ reveals a typical PL motif binding mode to VPS29 that superposes well with the previous cyclic peptide structures (**Figure 3D**) and other previous complexes with RidL, TBC1D5, VARP and VPS35L (38–42).

**Figure 3.**
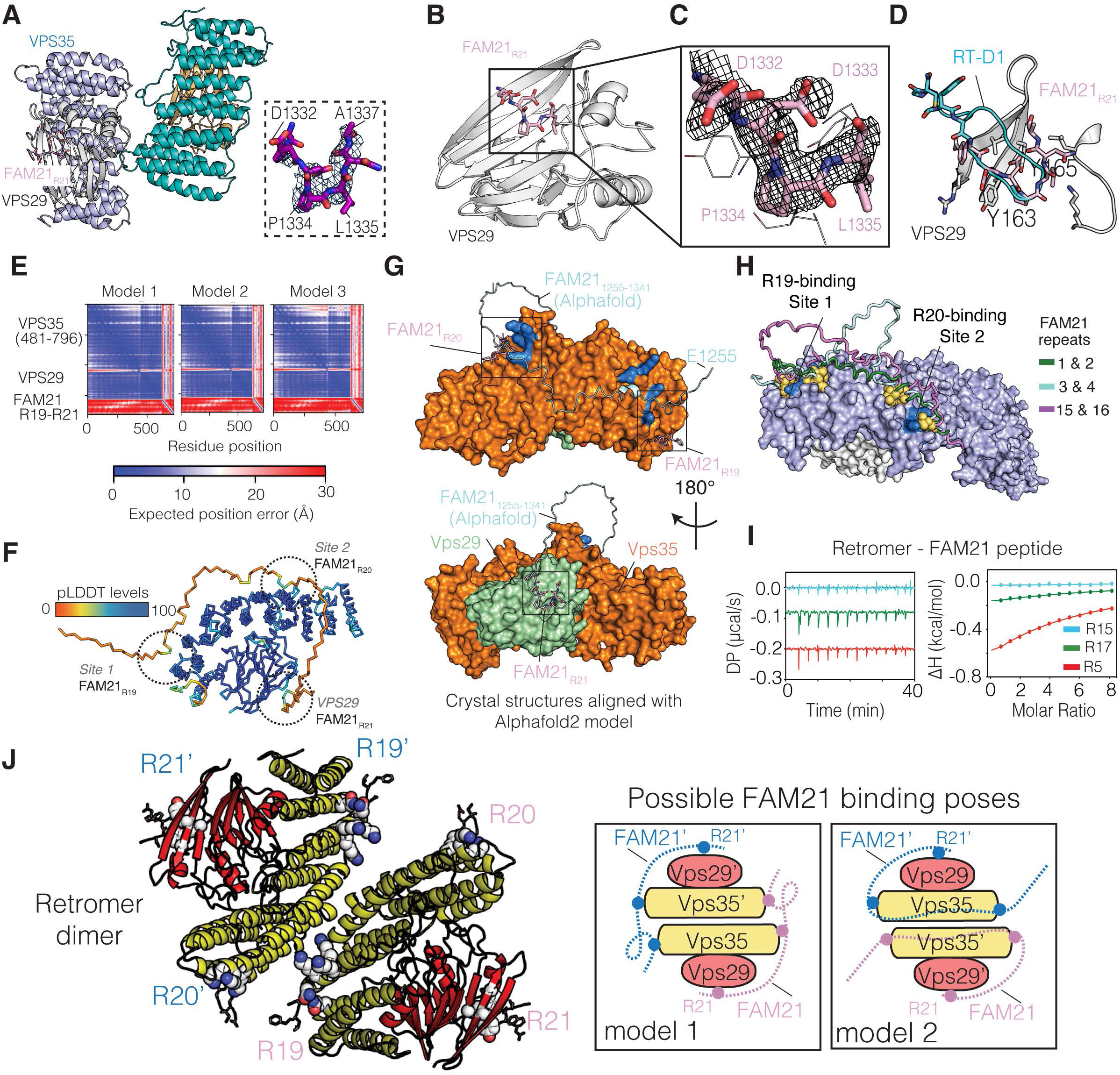
Structural basis for repeat 21 binding to VPS29 and model for Retromer engagement by multiple FAM21 repeats. (**A**) Crystal structure of VPS29–VPS35_481-796_ bound to the FAM21_R21_ peptide. (**B**) Crystal structure of VPS29 in complex with FAM21_R21_ peptide. (**C**) Enlarged view highlighting the refined density of FAM21_R21_ peptide peptide (2Fo–Fc map contoured at 1σ). (**D**) Structural alignment of the VPS29–FAM21_R21_ peptide complex with the VPS29–RT-D1 cyclic peptide complex (PDB ID 6XS5)(34) highlighting the PL motif binding region. (**E**) Predicted alignment Error (PAE) plots showing the predicted structure of VPS29–VPS35_481-796_–FAM21_R19-R21_ complex generated by Alphafold2. (**F**) The predicted structure highlighting the per-residue confidence score (pLDDT) between 0 to 100. Regions showing low pLDDT score (represents as green and orange) are likely to be unstructured. (**G**) Superimposition of all the two crystal structures with the Alphafold2 model. (**H**) Alphafold2 predictions of the complex of VPS29–VPS35 with either FAM21_R1-R2_, FAM21_R3-R4_ or FAM21_R15-R16_ showing pairs of repeats predicted to bind to the same positively charged grooves as repeats FAM21_19-20_. (**I**) ITC thermogram showing the binding between Retromer with FAM21_R5_ (red line), FAM21_R15_ (blue line) and FAM21_R17_ (green) peptides. All ITC graphs show the integrated (right) and normalized raw data (left) fit with a single-site binding model. (**J**) The locations of the FAM21 repeat 19 and 20 binding sites as well as the VPS29-binding repeat 21 are shown in the context of the VPS35-mediated Retromer dimer assembly derived from cryoelectron tomography (37) (PDB ID: 7BLN). The right panel shows two proposed binding models for how FAM21 can bind to a Retromer dimer in *cis* or *trans*.

Consistent with the two structures and our mutagenesis studies, AlphaFold2 predictions incorporating the FAM21 sequence with all three repeats 19-21 produced a model with the three repeats bound to the three distinct sites; R19 and R20 respectively binding Site 1 and Site 2 on VPS35 and R21 extending around to bind to the conserved VPS29 surface (**Figure 3E-3H**). Both FAM21_R20_ and FAM21_R21_ peptide observed in the crystal structures fit well to the various models generated by Alphafold2 (**Figure 3G**). It seems very likely other LFa repeats would have the capacity to interact with one or both sites in VPS35. Indeed further analysis using Alphafold2 supports the potential of other repeats to bind VPS35 with similar structures (**Figures 3H; Figure S3**), and we find that repeats 5 and 17 (although not repeat 15), show low affinity for Retromer (**Figure 3I**). Projecting these findings onto the VPS35-mediated Retromer dimer observed in cryotomography structures (37), we expect that the LFa repeats such as FAM21_R19,_ FAM21_R20_ and FAM21_R21_ can potentially use at least two distinct binding modes to engage the membrane-associated Retromer complex. With Repeat 21 bound to VPS29 as the reference point, repeats 19 and 20 and other LFa repeats from FAM21 could physically associate with two adjacent Retromer complexes in *trans* (model 1) or with one copy of Retromer in *cis* (model 2) (**Figure 3J**).

### Interaction of FAM21 aDLF motifs with the SNX27 FERM domain

In addition to Retromer, SNX27 has also been reported to bind FAM21 to enable proper cargo recycling (7, 43, 44). To confirm this interaction, we first examined the binding using co-immunoprecipitations with various truncation constructs of GFP-FAM21. Robust association was observed between SNX27 and C-terminal region of FAM21(1109-1341) and FAM21(1250-1341) (**Figure 4A**). A shorter construct of FAM21(1300-1341) however, which lacks all LFa repeats apart from repeat 21, was unable to bind. Using ITC, we found that full-length human SNX27 bound to GST-tagged FAM21_R19-21_ with a *K*_d_ of ∼6 μM (**Figure 4B; Table S3**). A similar binding affinity was also detected when the FERM domain (SNX27_FERM_) was used instead of the full-length SNX27 (**Figure 4B**), suggesting that the interaction primarily occurs through the FERM domain. We also tested the binding of SNX27_FL_ and SNX27_FERM_ with peptides corresponding to individual repeats FAM21_R19_, FAM21_R20_ and FAM21_R21_ (**Figure 4C and D**). The results further reinforced the co-immunoprecipitation experiments showing that both FAM21_R19_ and FAM21_R20_ peptides bind to SNX27 with a K_d_ of ∼7 μM and ∼20 μM respectively, while repeat 21 showed no detectable interaction.

**Figure 4.**
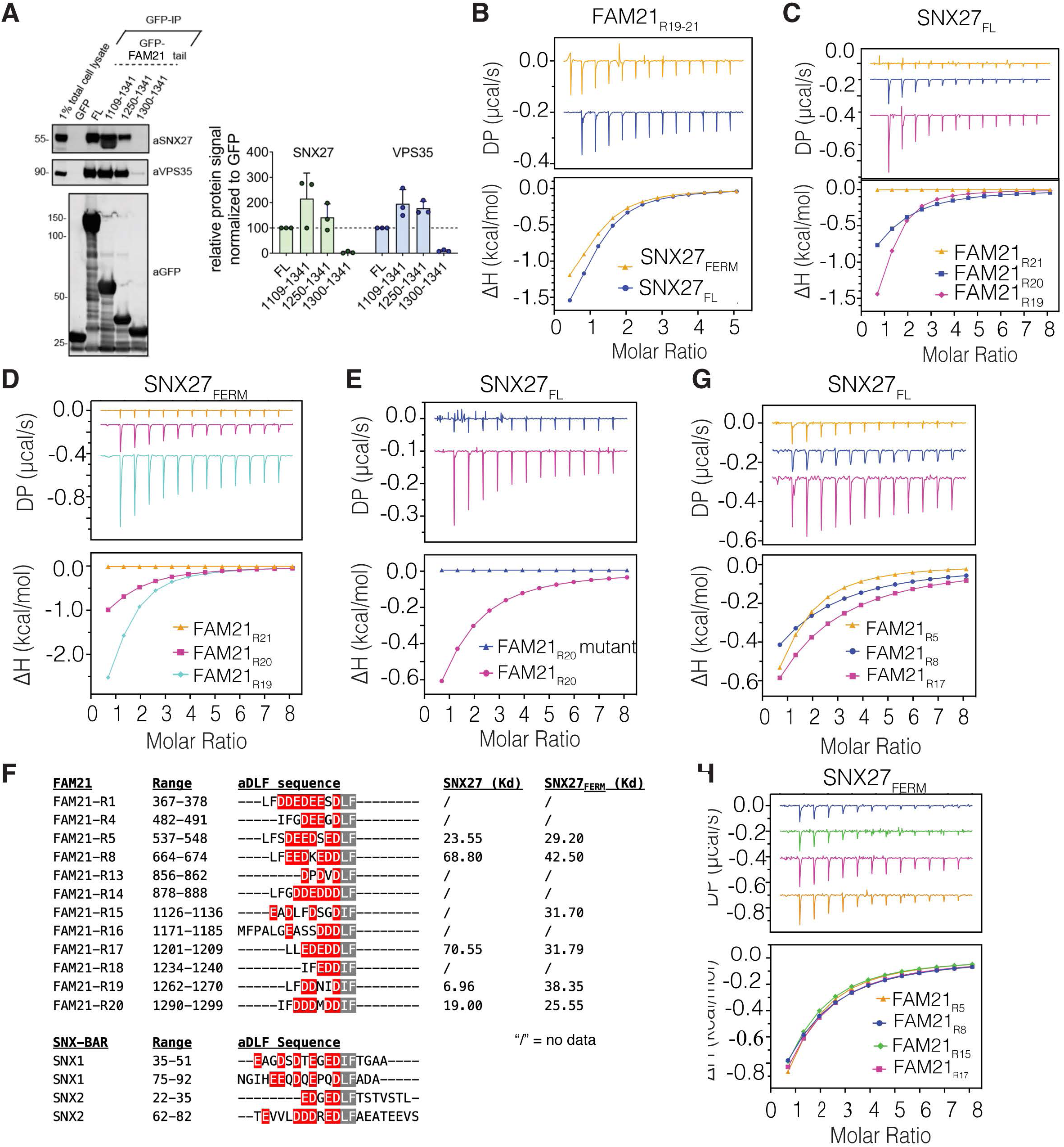
FAM21 aDLF repeats also binds to the SNX27 FERM domain. (**A**) Co-IP of GFP-tagged FAM21 tail in HEK293T cells. Samples were analysed by quantitative fluorescence-based western blotting. The band intensities of SNX27 and VPS35 were measured from n=3 independent experiments using Odyssey software. The SNX27 and VPS35 band intensities, normalized to GFP expression, are presented as the average fraction of the full length FAM21 tail. (**B**) ITC analysis showing the binding between GST-FAM21_R19-21_ and SNX27 (yellow line) and SNX27 FERM (blue line). (**C**) Similar binding experiment showing the binding of FAM21 peptides FAM21_R19_, FAM21_R20_ and FAM21_R21_ with either (**C**) SNX27 FL or (**D**) SNX27 FERM. (**E**) ITC thermogram comparing the binding between native FAM21_R20_ and ^1298^IF^1299^ to SS mutant peptides with SNX27 FL. (**F**) List of FAM21 peptides containing the same aDLF motif to those observed in SNX1 and SNX2. (**G)** ITC binding curves of the FAM21_R5_, FAM21_R8_ and FAM21_R17_ peptides titrated into either (**G**) SNX27 FL or (**H**) SNX27 FERM. All ITC graphs show the integrated (bottom) and normalized raw data (top) fit with a single-site binding model.

We noted that LFa repeats 19 and 20 (but not repeat 21) also contain sequences that resemble the acidic DLF (aDLF) motifs found in SNX1 and SNX2 previously shown to bind to the FERM domain of SNX27 (32). Indeed, mutations in the FAM21_R20_ peptide substituting the DIF sequence with DSS blocked interaction with SXN27_FL_ completely (**Figure 4E**), indicating that the D[L/I]F motif within repeats 19 and 20 play a major role in binding to SNX27. Sequence analysis of all of the 21 repeats within FAM21, showed that 12 of them also included the same sequence pattern with poly-acidic resides in front of a conserved D[L/I]F motif (**Figure 4F**). We proposed that all of these repeats are potentially able to bind SNX27, and using FAM21_R5_, FAM21_R8_, FAM21_R15_ and FAM21_R17_ peptides as examples we found all four showed micromolar binding affinity to hSXN27_FERM_ (**Figures 4G and 4H**). These ITC data together confirmed that the D[L/I]F motif is the main region responsible for binding to the FERM domain of SNX27, while the poly-acidic residues prior to the D[L/I]F motif play a critical role in facilitating the association.

### FAM21 and SNX1/SNX2 share the same binding pocket in the SNX27 FERM domain

To confirm the molecular basis for how FAM21 binds to SNX27, we first applied Alphafold2 to predict models of SNX27_FERM_ in complex with the subset of FAM21 repeats containing the putative aDLF motifs. Consistent with our expectation, all the FAM21 repeats were predicted to bind the same conserved pocket on the F3 module of the SNX27 FERM domain (**Figure S4**), also known to bind to the aDLF motifs from SNX1/SNX2 (32, 33). To validate the models we then co-crystallized SNX27_FERM_ with several different FAM21 repeat motifs and crystal structures were determined with FAM21_R15_, FAM21_R19_ and FAM21_R20_. All three complexes were solved in the same space group showing a single FERM domain in the asymmetric unit (**Figures 5A-5C; Table S4**). The electron density of the aDLF motif region was clearly visible in the highly conserved positively charged pocket of the SNX27 FERM domain (**Figures 5A-5C**). Superimposition of the three structures revealed a high degree of similarity without any major differences (**Figures 5D and 5E**). Our structures also superposed well to the SNX1 peptide bound complex published previously (32, 33) (**Figure 5F**).

**Figure 5.**
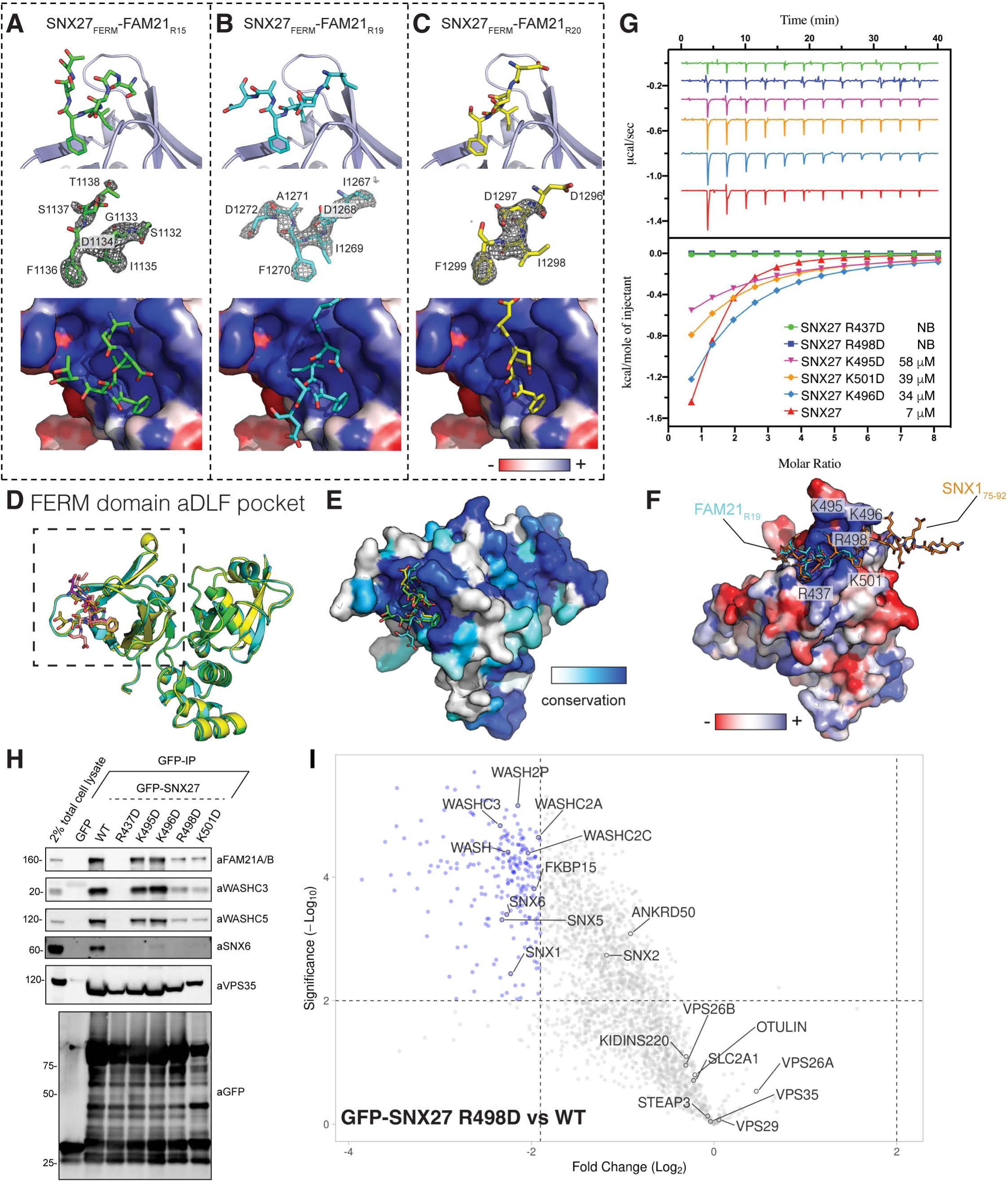
Structural basis for SNX27 interaction with FAM21 aDLF repeats. (**A**) Crystal structures of the SNX27 FERM domain bound to aDLF repeats of FAM21. Top views show the key region of SNX27 FERM involved in the binding to the FAM21 aDLF seqeunces (**A**) FAM21_R15_, (**B**) FAM21_R19_ and (**C**) FAM21_R20_. Middle panels show the electron density corresponding to a simulated-annealing composite omit 2Fo–Fc map contoured at 1σ. Bottom panels show electrostatic surface potential of SNX27 highlighting the positively charged pocket of SNX27 FERM. (**D**) Structural alignment showing the high degree of similarity between all three complex structures. (**E**) Sequence conservation map highlighting the conserved pocket of SNX27 FERM domain for binding to FAM21 aDLF motifs. (**F**) Structural alignment of SNX27 bound to FAM21_R19_ with the SNX27–SNX1_75-92_ complex (PDB ID 7CT1; (33)) showing they bind the same pocket. (**G**) ITC analysis showing the binding of the FAM21_R19_ peptide with the native or mutant forms of SNX27. The ITC graphs represent the integrated and normalized data fit with 1:1 ratio binding. (**H**) Co-IP of GFP-SNX27 mutants and binding of WASH, Retromer and ESCPE-1 (represented by SNX6) proteins assessed by Western blot. (**I**) Proteomic studies comparing the native SNX27 and mutated SNX27 R498D binding capabilities with SNX27 interactors assessed by Co-IP (n = 6; derived from 2 biological repeats each with 3 technical repeats). Mutation of the aDLF binding pocket causes disrupted interactions with most WASH and ESCPE-1 subunits. The full list of detected proteins is provided in **Dataset S1**.

To confirm the binding mode of SNX27 binding to FAM21 aDLF sequences we tested mutations in key SNX27 residues. SNX27 R437D and R498D mutants failed to interact with FAM21_R19_ peptide by ITC, while K495D, K496D and K501D reduced binding compared to the wild-type SNX27 (**Figure 5G**). The same pattern of binding was observed by co-immunoprecipitation of proteins from HEK293T cells (**Figure 5H**), although interestingly the K495D, K496D and K501D mutants had a greater impact on the ESCPE-1 complex interaction (represented by SNX6 in these experiments) than FAM21 and the WASH complex. This suggests that R437D and R498D play a central role in forming contacts with both FAM21 and SNX1/SNX2-derived aDLF peptides. In particular, R498 is likely to be responsible for electrostatic attraction with the poly-acidic residues in front of the aDLF motifs. Indeed, comparative unbiased proteomic analysis comparing the interactomes of wild-type SNX27 with the R498D mutant of SNX27 confirmed a significant reduction in association of SNX27(R498D) with subunits of both the WASH and ESCPE-1 complexes (**Figure 5I; Figure S5A; Table S2**).

### VPS35 Sites 1 and 2 enhance WASH endosomal recruitment but are not required for endosome-to-plasma membrane recycling of Retromer-SNX27 cargo proteins

We and others previously established that mutations in SNX27 that block interactions with aDLF motifs perturb endosomal retrieval and recycling of the SNX27-interacting cargo GLUT1 by the SNX27-Retromer-ESCPE-1 pathway (32, 33). While this phenotype was ascribed to a defect in SNX1 and SNX2-containing ESCPE-1 complex binding to SNX27 during the hand-over of retrieved cargo into newly forming tubular transport carriers (32), our results here indicate that the SNX27(R498D) mutant not only perturbs SNX1/SNX2 association but also binding to the WASH complex. Thus the defect in GLUT1 recycling could be attributed to either or both of these interactions, and it is not possible to design highly specific SNX27 mutations to distinquish them. Similarly, mutation of VPS29 to perturb the binding of FAM21 R21 will also preclude interactions with other essential proteins such as TBC1D5 and VARP (39, 45, 46). In addition, the highly redundant nature of the multiple overlapping LFa/aDLF repeats in FAM21 preclude the design of single point mutations in the FAM21 tail to test the role of specific sequences. Indeed, we found that although deletion of repeats 19-21 (residues 1250-1341) in the tail could reduce both Retromer and SNX27 interactions (**Figure 4A**), individual point mutations in these repeats had no impact on binding (**Figure S5B**) demonstrating that no single sequence is essential to mediate their binding.

To assess whether the interaction of the WASH complex with VPS35 was important for the endosomal activity we first assessed the localisation of FAM21 in VPS35 knock-out HeLa cells followed by rescue experiments with VPS35 mutations in both Site 1 and Site 2 (K555N/K556L/K559Q/K701E; NLQE mutant)(see **Figure 2G and 2H**). VPS35 KO cells show a modest but significant reduction in the colocalisation of FAM21 to SNX1-positive endosomal domains consistent with previous reports (12, 25). Re-expression of VPS35-GFP WT can partially rescue this mislocalistion, however VPS35(NLQE)-GFP is unable to restore FAM21 recruitment levels indicating the VPS35 interaction is important for promoting WASH complex localisation. Concistent with previous findings (47, 48), we did not observe any statistically significant alteration in the co-localization of CI-MPR with TGN46 in VPS35 KO cells, or in wild-type and mutant VPS35(NLQE) rescued cells **(Figure S5C**). We next assessed the importance of VPS35 interactions with FAM21 for receptor recycling. Rescue experiments were performed in both VPS35 knock-out HeLa cells (**Figure 6B and C**), and in H4 cells (**Figure S5D**) with similar results. As expected, in VPS35 KO cells the cell surface levels of SNX27-Retromer cargos including GLUT1 and Kidins220 were reduced. When GFP-VPS35 was re-expressed the cell surface levels of these proteins recovered partially or totally to normal levels. However, the VPS35(NLQE)-GFP mutant was also able to rescue endosomal recycling of these cargos, suggesting that reduced FAM21 binding to VPS35 on its own is not strictly required for the steady-state recycling of SNX27-Retromer-dependent cargos. Previous studies directly targeting FAM21 have shown that its depletion or knock-out leads to cargo sorting defects (43, 46). It is likely our data can be reconciled through the complex multivalent nature of FAM21 association with Retromer, such that although the VPS35 interaction with FAM21 reduces WASH interaction significantly (**Figure 2G and 2H**), and is required for a proportion of WASH recruitment to endosomal retrieval domains, the other minor sites in VPS29 retain an ability to engage FAM21 and may compensate for its loss. Further, FAM21 can interact with SNX27 and other endosomal proteins such as the Commander complex (25, 49), and the population of WASH associated with endosomes in the asbence of Retromer as seen here (**Figure 6A**) and in other studies (50) provide an added level of potential compensation.

**Figure 6.**
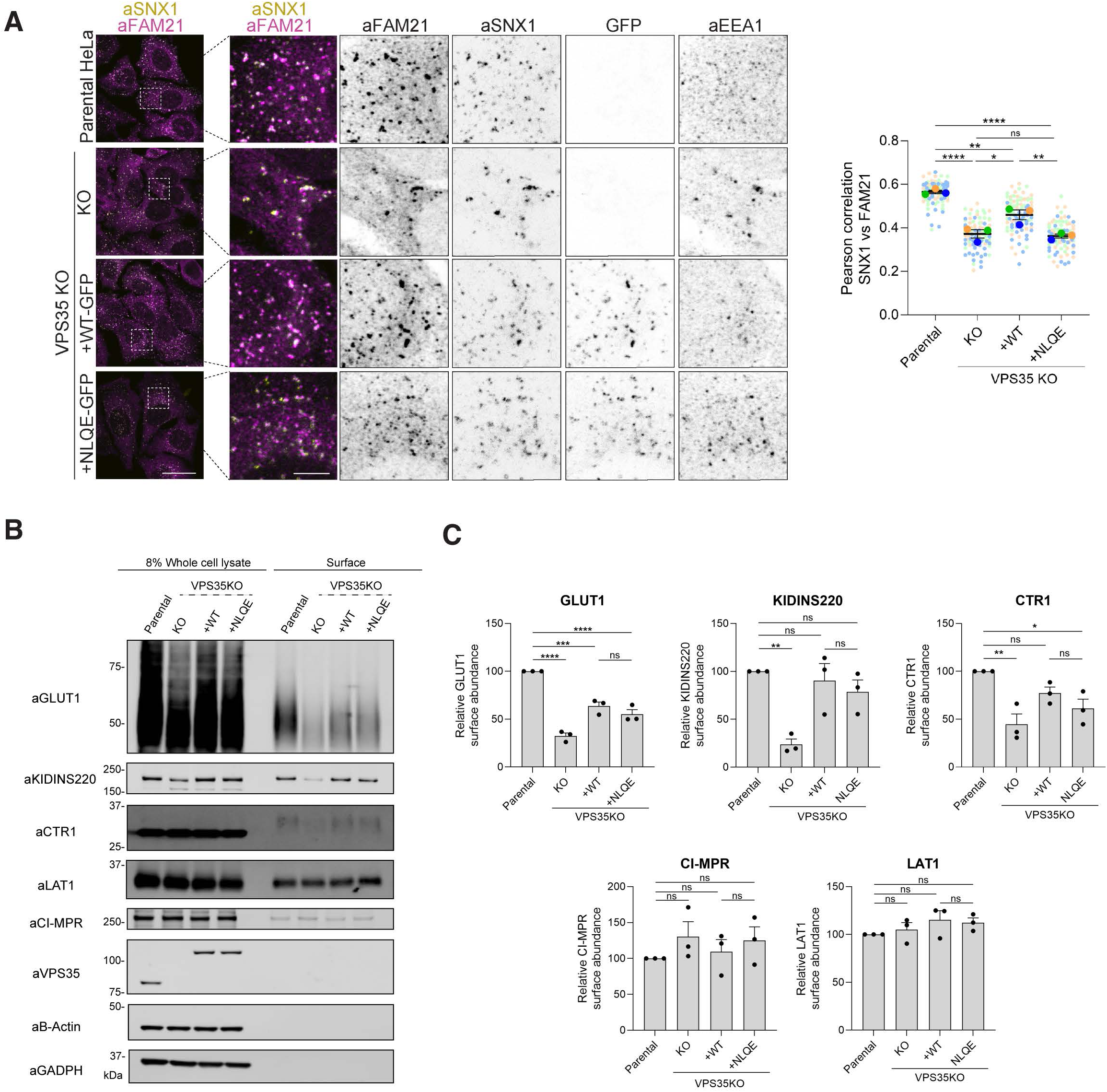
VPS35 interaction with FAM21 contributes to its endosomal localization, but it is not strictly required for SNX27–Retromer recycling. (**A**) VPS35 KO HeLa cells show a reduction in the colocalisation of FAM21 to SNX1-positive endosomal domains. Re-expression of VPS35-GFP WT can partially rescue this mislocalization, but the mutant VPS35(NLQE)-GFP with both FAM21 binding sites mutated is unable to restore FAM21 recruitment levels. A total of 90 cells were analyzed in each condition for colocalization between SNX1 and FAM21 across n=3 independent experiments. Scale bars, 25 μm (micrographs) and 5 μm (magnified images). (**B**) VPS35 KO HeLa cells showed reduced cell surface levels of SNX27–Retromer dependent cargos including GLUT1 and KIDINS220. VPS35 KO clonal line was stably transduced with lentiviral particles containing VPS35-GFP WT or VPS35(NLQE)-GFP. Re-expression and rescue with both constructs is able to rescue the cell surface levels of these cargos. This indicates that full binding of FAM21 to VPS35 is not strictly required for SNX27– Retromer mediated endosomal recycling. (**C**) Quantitation of Western blots is shown. Similar results were obtained in H4 cell lines as shown in **Figure S5C**. Statistics for western blots and confocal microscopy images were calculated from a minimum of 3 independent experimental repeats Graphs were plotted representing the mean value ± the standard error (SE) for each experimental condition. s. In all graphs, * = p < 0.05, ** = p < 0.01, *** = p < 0.001, **** = p < 0.0001.

## DISCUSSION

In this work we have defined the molecular mechanisms underpinning the direct interactions between Retromer, SNX27 and the WASH complex via the extended C-terminal domain of the FAM21 subunit. These interactions firmly establish the role of highly repetitive sequences in FAM21 for directly engaging both SNX27 and Retromer using distinct but overlapping sequence requirements. LFa repeats consist of two [L/I]F motifs interspersed by tracts of acidic side chains and mediate specific Retromer attachment through VPS35. Notably as this work was under review, a related study provided similar data relating to the Retromer interaction with FAM21 providing further support for the model (51). aDLF repeats consist of a D[L/I]F motif preceded by acidic residues, overlap with the LFa repeats and specifically interact with the FERM domain of SNX27 using a completely different mechanism. Combined with previous work (13, 26, 32), these studies lead to an evolving model for the assembly of the SNX27–Retromer–WASH endosomal recycling domain as one of extreme structural plasticity, sequence redundancy within FAM21, and molecular avidity between FAM21 and assembled oligomers of SNX27–Retromer on the endosomal membrane (**Figure S6**). This network of interactions will also incorporate other proteins known to interact with the FAM21 C-terminus, including the Commander assembly (25, 49, 52, 53), WDR91 (54), RME-8 (55), FKBP15 (13), as well as TBC1D23 which has also been shown to bind to FAM21’s LFa repeats (56, 57).

Our data shows that the LFa repeat sequences in FAM21 interact with Retromer at two distinct sites, and this is important for promoting WASH association with Retromer recycling domains, but not strictly essential for recycling of SNX27-Retromer cargos. Opposite to the VPS29-binding surface the VPS35 subunit has two separate sites for binding the peptide motifs, centred on K552, K555, K556 and K559 near the centre of the VPS35 α-helical solenoid (Site 2) and around K701, K705 and K749 near the C-terminus (Site 1). The discovery of two distinct binding pockets was surprising, and points to the potential of both redundancy in motif interactions (each site appears to have similar potential for binding FAM21 repeats) and avidity for increased binding strength (covalently connected sequences in FAM21 binding to two sites simultaneously). The presence of two binding sites is somewhat reminiscent of the binding of bipartite nuclear localisation signals to major and minor surfaces of the α-helical solenoid of importins (58), or of the nuclear receptors with the highly redundant FG repeats in the nucleoporins (59), although the mechanisms are completely different. Although we do not observe dramatic differences between the affinities of individual and multiple FAM21 repeats for Retromer by ITC, we propose that the combination of Retromer oligomerisation on the membrane (37, 60) with the multivalent nature of FAM21 interactions will promote a high level of specificity in WASH/Retromer engagement (61). Given the enormous extended structure of the FAM21 tail and the number of similar LFa motifs, FAM21 will readily interact with multiple Retromer complexes when organised into polymeric arches at the endosomal membrane. It is also worth noting that this will not lead to a specific stoichiometry for the Retromer/WASH interaction, but rather will promote formation of a heterogeneous and dynamic multivalent network.

In addition to the LFa sequences binding to VPS35, we identified a third binding site on the conserved surface of VPS29 for the final repeat of FAM21 which possesses a DPL sequence. This binds with modest affinity, and is likely inessential for direct Retromer-dependent endosomal recruitment of FAM21 (**Figure S5B**) (12, 13). We speculate however, that once the WASH complex has bound to Retromer and/or SNX27 that repeat 21 could then act to regulate the association of Retromer with other proteins that bind the same site such as TBC1D5 (39, 62, 63) or ANKRD27 (45) (**Figure S6**).

Retromer, SNX27 and WASH are intimately connected in the endosomal sorting and transport of numerous transmembrane proteins (64–66). The current model is that Retromer, along with SNX cargo adaptors such as SNX3 and SNX27, can establish endosomal retrieval sub-domains and membrane tubules enriched in cargo, and WASH is able to nucleate formation of branched actin filaments via Arp2/3 that enhance membrane tubulation and promote scission (8, 67). The capping-interacting motif (CPI) of FAM21 is responsible for recruiting uncapping dynactin mini-filaments, that then prime the WASH-induced actin branching (68). The first indications that Retromer and WASH cooperate in endosomal recycling demonstrated their colocalization and a direct interaction between the FAM21 C-terminal tail and the Retromer complex (10, 12, 13, 26). These studies also identified the 21 LFa repeats in the FAM21 tail and their importance for Retromer association (26). The C-terminal tail of FAM21 is sufficient for endosomal localisation (13), and Retromer is important for endosomal recruitment of WASH (12, 26, 27, 35, 36), although WASH can also be recruited through Retromer-independent mechanisms via FAM21 and WASHC4/SWIP subunits respectively (50). WASH depletion was also reported to result in exaggerated endosome-derived tubules (9, 10, 69), although this is not universally observed (43, 46, 70, 71).

Previous research established that the FERM domain of SNX27 associates with the aDLF motifs of SNX1 and SNX2 (32, 33), and our data now shows that FAM21 utilises an identical mechanism to engage with the SNX27 adaptor. As shown in **Figure 4F**, the sequence alignment with SNX1/SNX2 peptides shows the D[L/I]F triplet is the most important component for binding, and at least two acidic amino acids before this motif are also important for promoting electrostatic interactions with SNX27. SNX1 and SNX2 are SNX-BAR proteins and subunits of the ESCPE-1 complex along with SNX5 and SNX6 (32). Mutations targeting the aDLF motif in SNX1 or SNX2 that disrupted binding to SNX27 have highlighted the role of the SNX27-ESCPE-1 interaction in promoting the tubular-based endosomal exit and cell surface recycling of SNX27-Retromer cargo (32, 33). The results here both extend these previous analyses and complicate their interpretation, as the SNX27(R498D) FERM domain mutation disrupts not only ESCPE-1 binding but also FAM21 and WASH association. As it is impossible to design SNX27 mutations that only affect SNX1/SNX2 but not FAM21 binding (and *vice versa*), we currently cannot disentangle the individual roles of each interaction on SNX27-dependent trafficking, and effects could be due to either interaction or a combination of both. Overall, however, this further establishes that both Retromer and also SNX27 serve as direct ligands for overlapping multivalent repeat sequences in the FAM21 tail.

Combined with previous studies, our work makes it clear that the formation of endosomal retrieval sub-domains involves a dynamic network of generally low affinity interactions between Retromer, SNX27, ESCPE-1 and the WASH complex (and not discussed here ANKRD50) (44, 46), that includes a series of polyvalent extended sequences in subunits of the ESCPE-1 and WASH complexes. Low affinity interactions ensure that coincidence sensing of cargo and accessory proteins is required to provide the avidity necessary for pathway progression. One could consider these to be ‘checks and balances’ that derive from an instability in the protein interaction network when there is insufficient density of cargo and accessory proteins, then sub-domain organisation and coat formation will not be stabilised and the pathway will not progress (72). A more detailed understanding of both the spatial and temporal regulation of the interactions within the SNX27– Retromer–WASH–ESCPE-1 pathway will be challenging due to the high degree of multivalent and low affinity interactions involved, but will be essential for a complete portrait of this endosomal retrieval and recycling pathway(s). Reconstituting this pathway *in vitro* on artificial membranes, as well as high-resolution live-cell imaging will be required to begin to answer these questions, but the work here provides a key mechanistic foundation for these studies.

## Supporting information

Proteomic dataset

## ACKNOWLEDGEMENTS

We acknowledge use of the University of Queensland Remote Operation Crystallization and X-ray (UQ ROCX) Facility and the assistance of G.King and K.A.Arachchige. X-ray data were collected on the MX1 and MX2 microfocus beamline at the Australian Synchrotron.

## AUTHOR CONTRIBUTIONS

Initial Concept: B.M.C., and P.J.C.

Concept development: all authors.

ITC and structural determination and analysis: Q.G. and K.C.

Biochemistry and cell biology analysis: M.G-A., A.P.J., B.S., and C.M.D.

Proteomic analysis: K.J.H.

Data analysis and Figure generation: Q.G., K.C., A.P.J., B.S., and M.G-A.

Funding acquisition: B.M.C., and P.J.C.

Supervision: B.M.C., and P.J.C.

Writing – 1^st^ draft: Q.G. and K.C.

(Writing – review & editing: all authors)

## MATERIALS AND METHODS

Detailed methods are provided in the **Supporting Information Appendix**. DNA constructs, antibodies and other reagents used in this study are described in the **Supporting Information Appendix and Table S5.** All proteins used in this work were recombinantly expressed in *Eschericia coli* and purified using standard affinity chromatography and size exclusion chromatography. ITC experiments were performed using a MicroCal PEAQ ITC instrument operating at 25°C. Crystallographic X-ray diffraction data was collected at the Australian Synchrotron with structures determined by molecular replacement as outlined in **Table S2 and S4.** AlphaFold2 (73) predictions were performed using the open-source ColabFold pipeline (74). HeLa, HEK-293T and H4 cell lines were sourced from and authenticated by ATCC. Cells were grown under standard conditions in DMEM (Sigma-Aldrich) supplemented with 10% (v/v) FCS (Sigma-Aldrich) and 1% penicillin/streptomycin (Gibco, USA). Statistics for western blots and confocal microscopy images were calculated from a minimum of 3 independent experimental repeats.

