## Supporting Information Appendix for "Structural basis for coupling of the WASH subunit FAM21 with the endosomal SNX27-Retromer complex"

**This PDF file includes**
Supplementary Materials and Methods
Figures S1-S6
Tables S1-S5
Supplementary References

**SUPPLEMENTARY MATERIALS AND METHODS**

**MATERIALS AND METHODS:**

**Peptides**

All synthetic peptides applied for ITC and X-ray crystallography were obtained from Genscript (USA and Singapore). The peptides were freshly lyophilized using a laboratory freeze dryer Martin Christ ALPHA 1-2 LDplus to avoid chemical residues. To obtain the peptide stock solutions with a final concentration of 30 mM and a neutral pH, the freeze-dried peptides were dissolved in 1M HEPES pH 7.5. For use in ITC and co-crystallisation screening experiments, the peptides were diluted in protein buffer: 20 mM Tris-HCl pH 7.5, 200 mM NaCl, 5% glycerol, 1 mM β-ME. The sequences of peptides utilised were: FAM21-R5$_{536-552}$ GLFSDEEDSEDLFSSQS, FAM21-R8$_{662-678}$ TSLFEEDEEDDLFAIAK, FAM21-R15$_{1124-1140}$ RGEADLFDSGDIFSTGT, FAM21-R17$_{1197-1213}$ NPFPLLEDEDDLFTDQK, FAM21-R19$_{1261-1274}$ NLFDDNIDIFADLT, FAM21-R20$_{1289-1302}$ SIFDDDMDDIFSSG, FAM21-R20$_{1289-1302(IF/SS)}$ SSSDDDMDDSSSSG, and FAM21-R21$_{1328-1341}$ SNIFDDPLNAFGGQ, and they were generated from human WASH complex subunit 2A (FAM21A), Q641Q2.

**Molecular biology and cloning for recombinant protein production**

For bacteria expression of human Retromer and Retromer subcomplex, DNA encoding full-length Vps26A, Vps29, Vps35 and Vps35$_{481-796}$ were cloned into either pET28a or pGEX4T2 vectors as described previously (1). For SNX27, the codon-optimized gene containing full-length human SNX27 was synthesized by GenScript® and cloned into pET28a vector as described previously (2). The FERM domain containing residues 271 to 528 was inserted into pMCSG7 vector containing a N-terminal His-TEV tag. Human FAM21 truncation constructs including FAM21$_{R19-21}$, FAM21$_{R19-20}$, and FAM21$_{R20-21}$, were coned into pGEX4T2 vector for expression as a thrombin-cleavable N-terminal GST fusion protein. All the mutant constructs were conducted using site-directed mutagenesis approach with custom designed primers. All DNA constructs were verified using DNA sequencing.

**Protein expression and purification.**

All the protein constructs in this study were expressed using BL21 (DE3) competent cells and induced by isopropyl-β-D-1-thiogalactoside (IPTG) approach. In all cases unless noted otherwise, the cells were cultivated in Ultra Yield™ flasks at 37˚C and 180 rpm until the cell density reached $OD_{600}$ ~0.8. A final concentration of 0.8 mM IPTG was then added to the expression culture and further incubated for 16 h to 18 h at 20˚C. To achieve the optimal expression yield of hSNX27$_{FERM}$, a final concentration of 0.5 mM IPTG was added to the culture prior to the incubation at 20˚C. After expression, the cells were pelleted and harvested by 10 min of centrifugation using JLA 8.1 rotor at 6000 rpm, 4˚C. The cell pellets were either stored at -80˚C or immediately proceeded to purification.

Purification of Retromer was performed using the protocol described previously (1). In brief, the GST-tagged VPS29 co-expressed with VPS35 were mixed with the cell pellet of VPS26A. Cells were lysed through a Constant System TS-series high pressure cell disruptor in lysis buffer containing 50 mM Tris-HCl pH 7.5, 200 mM NaCl, 2 mM 2-Mercaptoethanol (β-ME), 1 mM benzamidine, 10 μg/mL deoxyribonuclease I (DNase I). The soluble homogenate cleared by centrifugation was loaded onto Talon resin (Clonetech) followed by glutathione Sepharose (GE Healthcare) to obtain the correct stoichiometry ratio of the Retromer complex. For the VPS29 – $VPS35_{481-796}$ subcomplex and VPS29 alone, the purification was carried out using glutathione Sepharose (GE Healthcare) directly. Removal of the fusion tag from Retromer, subcomplex and VPS29 were performed using on-column cleavage with thrombin protease (Sigma-Aldrich) overnight at 4˚C. The flow-through containing the fusion tag free protein was further purified using size-exclusion chromatography (SEC) on a HiLoad® 16/600 Superdex 200 or Superdex 75 column equilibrated with buffer containing 50 mM Tris-HCl pH 7.5, 200 mM NaCl, 2 mM β-ME. Purification of hSNX27 constructs were carried our using protocol described previously (2). For the FAM21 constructs, the cell pellet of GST-tagged FAM21 protein was resuspended in lysis buffer containing 20 mM Tris-HCl pH 7.5, 200 mM NaCl, 5% glycerol, 2 mM β-ME, 1 mM benzamidine, and 10 μg/mL DNase I. Cells were lysed and centrifuged using the same protocol as Retromer and SNX27. Supernatants of GST-tagged protein homogenates were loaded onto glutathione Sepharose$^{TM}$ beads (GE healthcare). Prior to the elution, the protein-bound bead was washed with 100 ml of washing buffer containing 20 mM Tris-HCl pH 7.5, 200 mM NaCl, 2 mM β-ME. The elution buffer then carried out using the same buffer as the washing buffer supplemented with freshly prepared 20 mM reduced glutathione and the 1 X cOmplete$^{TM}$ EDTA-free protease inhibitor tablet. The protease inhibitor table was added to protect the GST-tag from non-specific cleavage during purification. The SEC was performed as the final purification step using HiLoad® 16/600 Superdex 200 or Superdex 75 (GE Healthcare) column pre-equilibrated with buffer containing 50 mM Tris-HCl pH 7.5, 200 mM NaCl, 2 mM β-ME. Protein integrity was validated using SDS-PAGE gel and concentration was measured using Nanodrop at $OD_{280.}$

**Isothermal titration calorimetry (ITC)**

All microcalorimetry experiments were carried out at 25˚C using a PEAQ ITC (Malvern, UK) in SEC buffer. The interaction of Retromer and FAM21 peptides was carried out by titrating 400 - 600 μM of FAM21 peptides into 15 - 20 μM of Retromer, VPS29 – $VPS35_{481-796}$ subcomplex or VPS29. The effect of the cyclic peptide was performed by titrating 400 μM of FAM21 R21 peptides into 15 μM of VPS29 supplemented with 2 mM RT-D1. In the native control, the cyclic peptide was substituted with equivalent percentage (v/v) of DMSO. For the Retromer mutant experiment, 400 – 600 μM of GST-$FAM21_{R19-R20}$ or 600 μM of FAM21 R21 peptide was titrated into either 15 - 20 μM of native Retromer or $Retromer_{K555N,\ K556L,\ K559Q}$ mutant. In the case of SNX27 and FAM21 interaction, 300 μM of GST-$FAM21_{R19-21}$ was titrated into 12 μM of either $hSNX27_{FL}$ or $hSNX27_{FERM}$. For the binding tests of FAM21 peptides to SNX27, 800 μM of the FAM21 peptides were titrated into 20 μM $hSNX27_{FL}$ or $hSNX27_{FERM}$ proteins. Mutant binding experiment was performed by titrating 600 μM of $FAM21_{R19}$ peptide into 20 μM of either native or single point mutant form of $hSNX27_{FL}$.

All ITC experiments were performed with a single injection of 0.4 μL followed by a series of 12 injections of 3.2 μL each, spaced 180 seconds apart, and stirred at 850 rpm. By integrating the observed peaks and subtracting the heats of dilution from the background, the heat exchange of interactions was calculated. The thermodynamic parameters Kd, ΔH, ΔG, and −TΔS for all binding tests were obtained by fitting and normalising data to a single-site binding model using the Malvern software package. The stoichiometry was adjusted initially, and if the value was nearly 1, N was set to exactly 1.0 for computation. To ensure the data were reproducible, each experiment was run at least twice.

**Crystallisation**

Crystallization screening was carried out using the hanging-drop vapour diffusion method at a temperature of 20˚C. To obtain the FAM21 R20 bound co-crystal structure, the purified VPS29 – $VPS35_{481-796}$ subcomplex was mixed with 8 molar excesses of $FAM21_{R20}$ peptide solubilized in 1 M HEPES pH 7.5. Crystals were observed in multiple different commercial screen conditions. Best diffracting quality crystals were obtained using streak seeding approach in 24-well plate with a final protein concentration of 10 mg/ml under condition comprising 0.1 M Sodium Citrate pH 4.2, 0.2 M NaCl 16% PEG1000. Rod shape crystals appeared after 1 day and grew to optimal after 5 days incubation at 20˚C. In the case of FAM21 R21 bound co-crystal structure, two different protein complexes were prepared. In the first case, the purified VPS29 – $VPS35_{481-796}$ subcomplex was mixed with 5 molar excesses of $FAM21_{R21}$ peptide solubilized in 1 M HEPES pH 7.5 at a final concentration of 12 mg/ml in condition containing 0.1 M MES pH 6.5, 0.1 M Magnesium Acetate, 10% PEG10,000. Crystals appeared after 3 days of incubation at 20˚C. In the second case, the purified VPS29 was mixed with 2 molar excesses of $FAM21_{R21}$ peptide at a final protein concentration of 15 mg/ml. The protein-$FAM21_{R21}$ peptide complex crystals were obtained in condition composed of 0.2 M Magnesium Formate, 20% PEG3350. Crystals appeared after 2 weeks of incubation at 20˚C.

To co-crystallize $hSNX27_{FERM}$ with FAM21 peptides, five-fold molar excesses of the $FAM21_{R15}$, $FAM21_{R19}$ and $FAM21_{R20}$ peptides were added respectively to the purified $hSNX27_{FERM}$ to a final concentration of 17 mg/ml. Additionally, 10 mM DTT was supplemented to protect free cysteines. For co-crystallisation with peptide $FAM21_{R20}$, $hSNX27_{FERM}$ was buffer exchanged to buffer containing 20 mM Tris pH 7.5, 100 mM NaCl, 5% glycerol and 1 mM β-ME. Initial needle-shaped crystals were observed in the commercial screen condition ProPlex C11, which contains 0.1 M Sodium Cacodylate pH 6.5 and 25% PEG4000. The best diffraction-quality crystals were produced in 24-well plate under conditions comprising 0.1 M Sodium Cacodylate pH 6.0 and 22.5% PEG4000; 0.1 M Sodium Cacodylate pH 6.5 and 25% PEG3350; 0.1 M Sodium Cacodylate pH 6.5 and 22.5% PEG4000 for $hSNX27_{FERM}$-$FAM21_{R15}$, $hSNX27_{FERM}$-$FAM21_{R19}$ and $hSNX27_{FERM}$-$FAM21_{R20}$ complexes respectively.

**Crystallographic data collection and structure determination**

The X-ray diffraction data were collected at 100 K on the MX2 beamlines at the Australian Synchrotron. All the collected data were indexed and integrated by AutoXDS (3) and scaled using Aimless (4). Phase was solved by molecular replacement using Phaser (5). Crystal structures of VPS29 – $VPS35_{481-796}$ – $FAM21_{R20}$ and VPS29 – $VPS35_{481-796}$ – $FAM21_{R21}$ were solved using Retromer subcomplex hVPS29 – $hVPS35_{483-780}$ (PDB ID: 2R17) as the initial model. In the case of VPS29 – $FAM21_{R21}$ complex, human VPS29 from Vps29 – RT-D2 complex structure (PDB ID: 6XS7) was used as the template. For $hSNX27_{FERM}$ – $FAM21_{R15}$, $hSNX27_{FERM}$ – $FAM21_{R19}$ and $hSNX27_{FERM}$ – $FAM21_{R20}$ complexes, the $hSNX27_{FERM}$ model generated by AlphaFold2 was used as the template. Structure refinement was performed using the PHENIX suite (6) with iterative rebuilding of the model. The refined model was manually rebuilded using Coot guided by Fo - Fc difference maps. Data collection and refinement statistics are summarized in **Supplementary Table S3**. Sequence conservation was based on a T-Coffee multiple sequence alignment (7) and calculated using the ConSurf Server (8). Structural alignments and molecular figures were generated using PyMOL (Schrodinger, USA).

**AlphaFold predictions of FAM21 peptides binding to $hSNX27_{FERM}$**

We utilised the AlphaFold2 neural network (9) of the open-source ColabFold pipeline (10) to produce models of FAM21 peptides that associate with Retromer or the SNX27 FERM domain. In the case of Retromer – FAM21 complex, the sequences of VPS29, VPS35 and FAM21 repeats 1, 2, 3, 4, 15, 16, 19, 20 and 21 were applied. For SNX27 – FAM21 complex, 12 of the acidic DLF motifs of the FAM21 were modelled with the sequence of the human SNX27 FERM domain (residues 271 to 528, NP_112180). The default settings for ColabFold were used to construct multiple sequence alignments using MMseqs2 (11) and perform structural relaxing of the final peptide geometry with Amber (12), which resulted in the generation of three models for each peptide. Structural alignments and molecular figures were generated using PyMOL (Schrodinger, USA).

**Antibodies**

Antibodies used in this study were mouse monoclonal antibodies: GFP (clones 7.1 and 13.1; 11814460001; Roche, Germany) (1:2,000 for WB), DMT-1(sc-166884, Santa Cruz Biotechnology) (1:1,000 for WB), GADPH (97166, Cell Signaling Technologies) (1:1,000 for WB), ITGb1 (clone 18, BD610467, BD Biosciences) (1:1,000 for WB), N-Cadherin (13A9, Cell signaling technology) (1:1,000 for WB), SNX1 (clone 51/SNX1; 611482; BD, USA) (1:1,000 for WB, 1:200 for IF), SNX2 (clone 13/SNX2; 5345661; BD) (1:1,000 for WB), SNX6 (clone d-5, 365965; Santa Cruz Biotechnology, USA) (1:1,000 for WB), SNX27 (ab77799, Abcam, UK) (1:1,000 for WB), β-actin (A1978; Sigma-Aldrich, USA) (1:2,000 for WB), VPS29 (clone d-1, 398874; Santa Cruz Biotechnology) (1:200 for WB), WASHC5 (clone B-10, sc-377146) (1:1,000 for WB); rabbit monoclonal antibodies: CI-MPR (EPR6599; 124767; Abcam) (1:1,000 for WB, 1:200 for IF), VPS35 (EPR11501(B); 157220; Abcam) (1:1,000 for WB), GLUT1 (ab115730, Abcam) (1:1,000 for WB); rabbit polyclonal antibodies: FAM21 (kind gift from Dan Billadeau) (1:1000 for WB, 1:400 for IF), KIDINS-220 (21856-1-AP, Proteintech) (1:1,000 for WB), LAT1 (5347S, Cell signaling technology) (1:1,000 for WB), GFP (GTX20290; GeneTex, USA) (WB 1:2,000), STEAP-3 (17186-1-AP, Proteintech) (1:1,000 for WB), VPS26A (23892; Abcam) (1:1,000 for WB), VPS35 (97545; Abcam) (1:1,000 for WB), VPS29 (98929; Abcam) (1:100 for WB), WASH1 (kind gift from Dan Billadeau) (1:1,000 for WB), WASHC4 (ABT69, Merck) (1:1,000 for WB); goat polyclonal antibody EEA1 (A121550, Antibodies.com) (1:200 for IF); sheep polyclonal antibody TGN46 (GTX74290, GeneTex) (1:200 for IF).

**Cell culture and transfection**

HeLa, HEK-293T and H4 cell lines were sourced from and authenticated by ATCC. Cells were grown under standard conditions in DMEM (Sigma-Aldrich) supplemented with 10% (v/v) FCS (Sigma-Aldrich) and 1% penicillin/streptomycin (Gibco, USA). For GFP-based immunoprecipitations, HEK293T cells were transfected with GFP constructs using polyethylenimine (Sigma-Aldrich) and expression was allowed for 24-48 hours. To produce stably expressing cells, HeLa and H4 cells were transduced with lentiviruses with the constructs cloned in pXLG3.

**Biotinylation of cell surface proteins**

For surface biotinylation experiments, fresh Sulfo-NHS-SS Biotin (Thermo Fisher Scientific, #21217) was dissolved in ice-cold PBS at pH 7.8 at a final concentration of 0.2 mg/ml. Cells were washed twice in ice-cold PBS and placed on ice to slow down the endocytic pathway. Next, cells were incubated with the biotinylation reagent for 30 minutes at 4˚C followed by incubation in TBS for 10 minutes to quench the unbound biotin. The cells were then lysed in lysis buffer and equal amounts of their proteins were subjected to Streptavidin beads-based affinity isolation (GE Healthcare, USA). After 30 minutes incubation at 4 degrees, the beads where washed 3 times with PBS + 1% TX100 and 1.2M NaCl.

**Immunoprecipitation and quantitative western blot analysis**

For western blotting, cells were lysed in PBS with 1% (v/v) Triton X-100 and Pierce protease inhibitor cocktail (Thermo Scientific). The protein concentration was determined with a BCA assay kit (Thermo Fisher Scientific, USA), and equal amounts were resolved on NuPAGE 4% to 12% precast gels (Invitrogen, USA). Blotting was performed onto polyvinylidene difluoride membranes (Immobilon-FL; EMD Millipore, USA) followed by detection using the Odyssey infrared scanning system (LI-COR Biosciences, USA). For GFP-based immunoprecipitations, HEK-293T cells were lysed 48 hours after transfection in immunoprecipitation buffer (50 mM Tris-HCl, 0.5% (v/v) NP-40, and Roche protease inhibitor cocktail) and subjected to GFP trap (ChromoTek, Germany). Immunoblotting was performed using standard procedures. Detection was performed on an Odyssey infrared scanning system (LI-COR Biosciences) using fluorescently labeled secondary antibodies.

**TMT-labelling, Nano LC-MS/MS and Proteomic data analysis**

Were performed as described previously (13, 14).

**Immunofluorescence staining, image acquisition and analysis**

HeLa cells were seeded onto 13mm coverslips. One day after, cells were washed in PBS, fixed in 4% (v/v) for 20 minutes and washed again 3 times. Cells were permeabilized with 0.1% (v/v) Triton X-100, blocked in 1%

(w/v) BSA, incubated for 1h with primary antibodies followed by 30 minutes with secondary antibodies (Alexa Fluor, Thermo Fisher Scientific) in 1% BSA. Coverslips were mounted onto glass microscope slides with Fluoromount-G (Invitrogen). Microscopy images were collected at 37 °C with a Leica SP8 AOBS confocal laser scanning microscope attached to a Leica DM I8 inverted epifluorescence microscope (Leica Microsystems, Germany), with a 63x 1.4 NA oil immersion lens (506350, Leica Microsystems). Images were acquired using LASX software (Leica microsystems). Colocalisation and fluorescence intensity analysis were performed using Volocity 6.3 software (PerkinElmer) with automatic Costes background thresholding. Immunofluorescence images were prepared in Volocity 6.3, ImageJ and Adobe Illustrator (Adobe).

**Statistical analysis of western blots and confocal microscopy**

Statistics for western blots and confocal microscopy images were calculated from a minimum of 3 independent experimental repeats. Graphs were generated using GraphPad Prism 9 software (LaJolla, CA) plotting the the mean value ± the standard error (SE) for each experimental condition. n represents the number of independent experimental repeats. One-way ANOVA with Dunnett's multiple comparison test was used to assess statistically significance. In all graphs, * = $p < 0.05$, ** = $p < 0.01$, *** = $p < 0.001$, **** = $p < 0.0001$.

### SUPPLEMENTARY FIGURES AND DATA

**Figure S1. Sequence alignment of FAM21/WASHC2 repeats 19, 20 and 21.**

Repeats 19-21 of FAM21 are aligned from various species. Conserved [I/L]F dipeptide motifs are indicated by yellow arrows while the specific PL motif in repeat 21 is indivated by green ovals. h, homo sapiens; m, Mus musculus; r, Rattus norvegicus; b, Bos taurus; d, Drosophila melanogaster; zf, Danio rerio.

**Figure S2. Quantitation of the Western blots shown in Figure 2G and 2H.**

Band intensities from **Figure 2G and 2H** were quantified using Odyssey software and normalized to GFP expression. 1-way ANOVA with Dunnett's multiple comparisons test. Error bars represent the standard error. $^*P < 0.05$, $^{**}P < 0.01$, $^{***}P < 0.001$, $^{****}P < 0.0001$.

**Figure S3. Alphafold models of VPS35–VPS29 in complex with different FAM21 LFa repeats.**

(**A**) Predicted alignment Error (PAE) plots for the AlphaFold2-predicted structures of the human (**A**) VPS35–VPS29–FAM21$_{R1\text{-}R2}$ complex, (**B**) VPS35–VPS29–FAM21$_{R3\text{-}R4}$ complex, (**C**) VPS35–VPS29–FAM21$_{R15\text{-}R16}$ complex. On the right the predicted structures are shown coloured according to the per-residue confidence score (pLDDT) between 0 to 100. Regions showing a low pLDDT score (green and orange) are likely to be unstructured.

**Figure S4. Alphafold models of SNX27 FERM domain in complex with various FAM21 aDLF sequences.**

(**A**) Overlay of all AlphaFold2 predicted structures shown as Cα backbone ribbon. The sequences of the aDLF repeats are shown in **Figure 4F.** (**B**) The predicted complex of SNX27$_{FERM}$ bound to FAM21 aDLF sequence from repeat 1. SNX27 is shown in cartoon representation coloured according to pLDDT score. The FAM21$_{R1}$ peptide is shown in green sticks. (**C**) Predicted alignment Error (PAE) plots showing the predicted structure of human SNX27$_{FERM}$–FAM21 aDLF complexes. The top ranked prediction is shown in close up in the left hand panels.

**Figure S5. Interactions of SNX27 and FAM21 mutants and VPS35 rescue experiments in H4 cells.**

(**A**) Co-IP in HEKs of GFP-SNX27 wild-type and R498D mutant and binding of WASH, Retromer (indicated by subunit VPS26A) and model cargo KIDINS220 assessed by Western blot. (**B**) Co-IP of GFP-FAM21 and mutants shows that specific mutations in repeats 19-21 do not prevent binding to Retromer or SNX27 in cells. KK is the combination of D1268K and D1297K mutants. (**C**) CIMPR colocalization with TGN46 is not altered in VPS35 KO, wild-type or NLQE mutant rescued HeLa cells. A total of 70 cells were analyzed in each condition across n=3 independent experiments. Scale bars, 25 µm (micrographs) and 5 µm (magnified images). (**D**) VPS35 KO H4 cells showed reduced cell surface levels of SNX27–Retromer dependent cargos including GLUT1 and KIDINS220. Re-expression and rescue with both GFP-VPS35 wild-type protein and VPS35(NLQE)-GFP is able to partially rescue the cell surface levels of these cargos. Band intensities for all blots were quantified using Odyssey software and normalized to GFP expression. 1-way ANOVA with

Dunnett's multiple comparisons test. Error bars represent the mean, s.d. *$P < 0.05$, **$P < 0.01$, ***$P < 0.001$, ****$P < 0.0001$.

**Figure S6. Summary and proposed model for the cooperative recruitment of FAM21 by Retromer, SNX27 and other endosomal proteins.**

The WASH subunit FAM21 has 21 LFa repeated elements in its extended tail that are rich in acidic, isoleucine and phenylalanine residues. LFa sequences possess two [L/I]F motifs interspersed by an acidic stretch ([L/I]F$[D/E]_{3\text{-}9}$[L/I]F). These bind two conserved sites towards the C-terminus of the VPS35 subunit of Retromer. The last repeated element of FAM21 has a specific PL dipeptide sequence that promotes interaction with Retromer subunit VPS29. This would be mutually exclusive of binding to other VPS29 interactors such as TBC1D5, ANKRD27, VPS35L and bacterial effector RidL. aDLF sequences are a subset of the LFa repeats with an acidic stretch prior to a D[L/I]F motif that associate with a conserved pocket in the SNX27 FERM domain, and are also found in the SNX1 and SNX2 proteins of the ESCPE-1 complex.

**Dataset S1. Proteomic interactions of GFP-SNX27(R498D) mutant versus GFP-SNX27 wild type.**

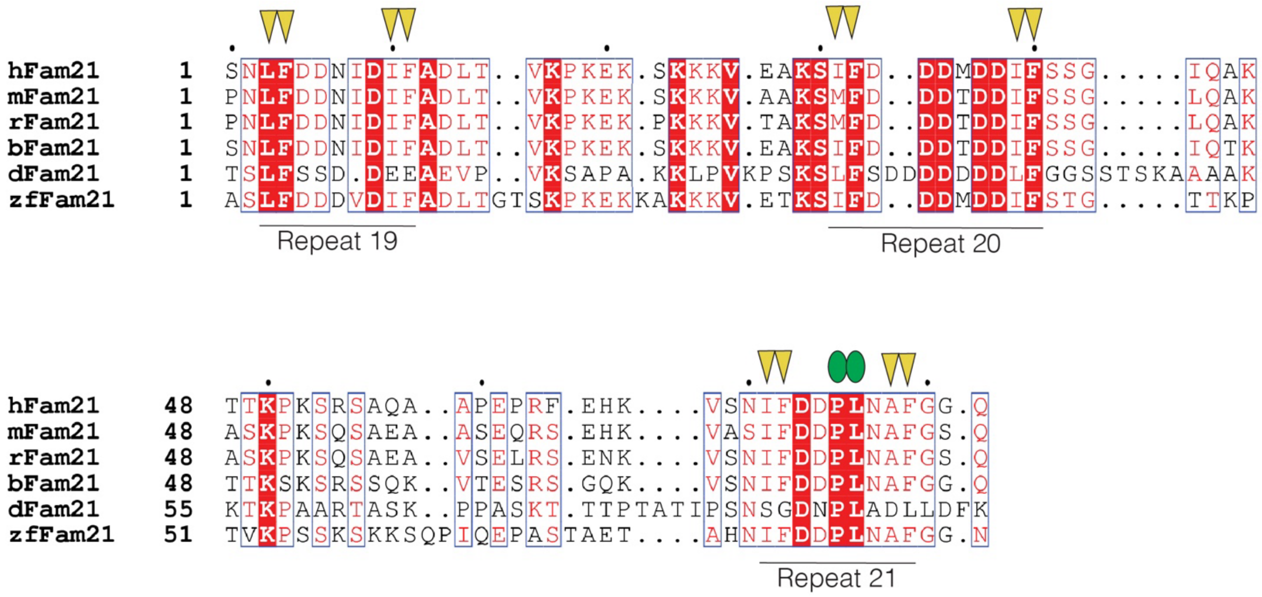

**Figure S1. Sequence alignment of FAM21/WASHC2 repeats 19, 20 and 21.**
Repeats 19-21 of FAM21 are aligned from various species. Conserved [I/L]F dipeptide motifs are indicated by yellow arrows while the specific PL motif in repeat 21 is indivated by green ovals. h, homo sapiens; m, Mus musculus; r, Rattus norvegicus; b, Bos taurus; d, Drosophila melanogaster; zf, Danio rerio.

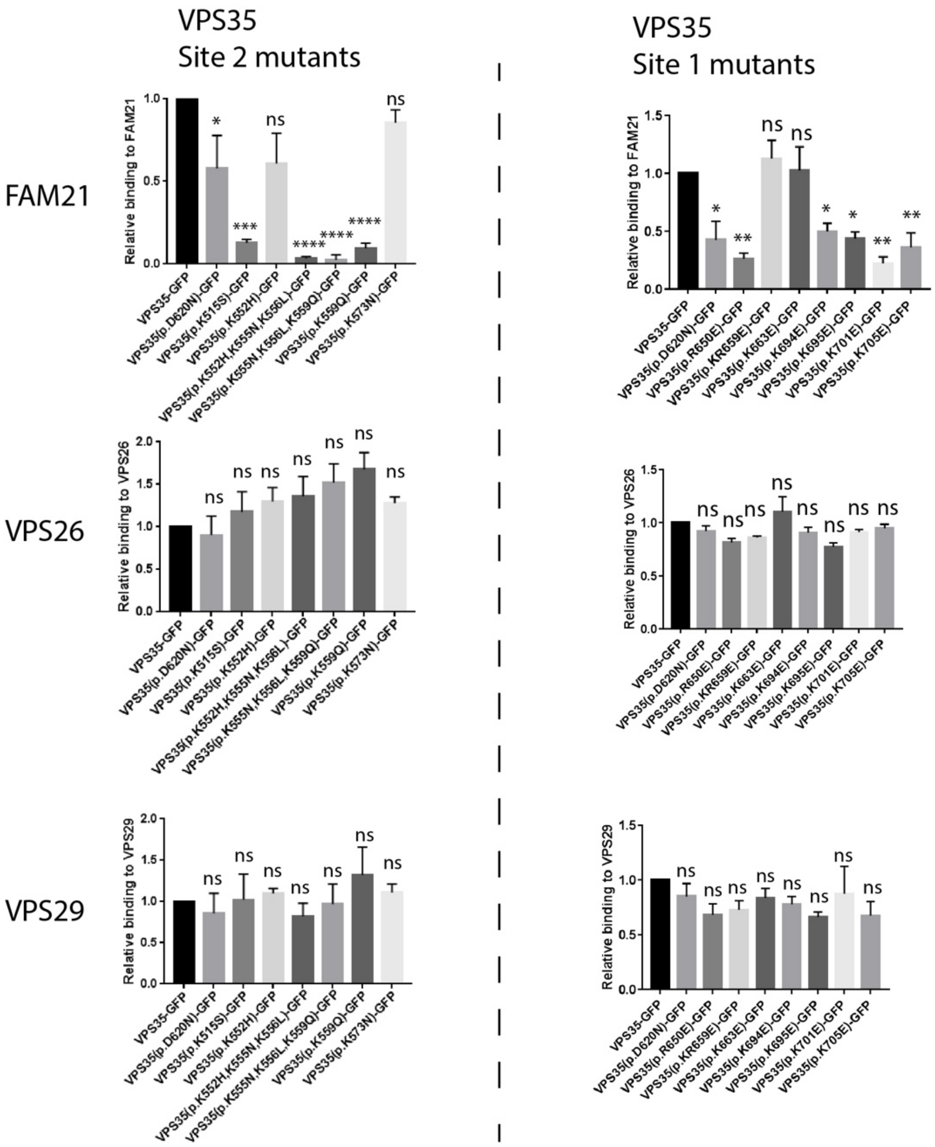


**Figure S2. Quantitation of the Western blots shown in Figure 2G and 2H.**

Band intensities from Figure 2G and 2H were quantified using Odyssey software and normalized to GFP expression. 1-way ANOVA with Dunnett's multiple comparisons test. Error bars represent the standard error. *P < 0.05, **P < 0.01, ***P < 0.001, ****P < 0.0001.

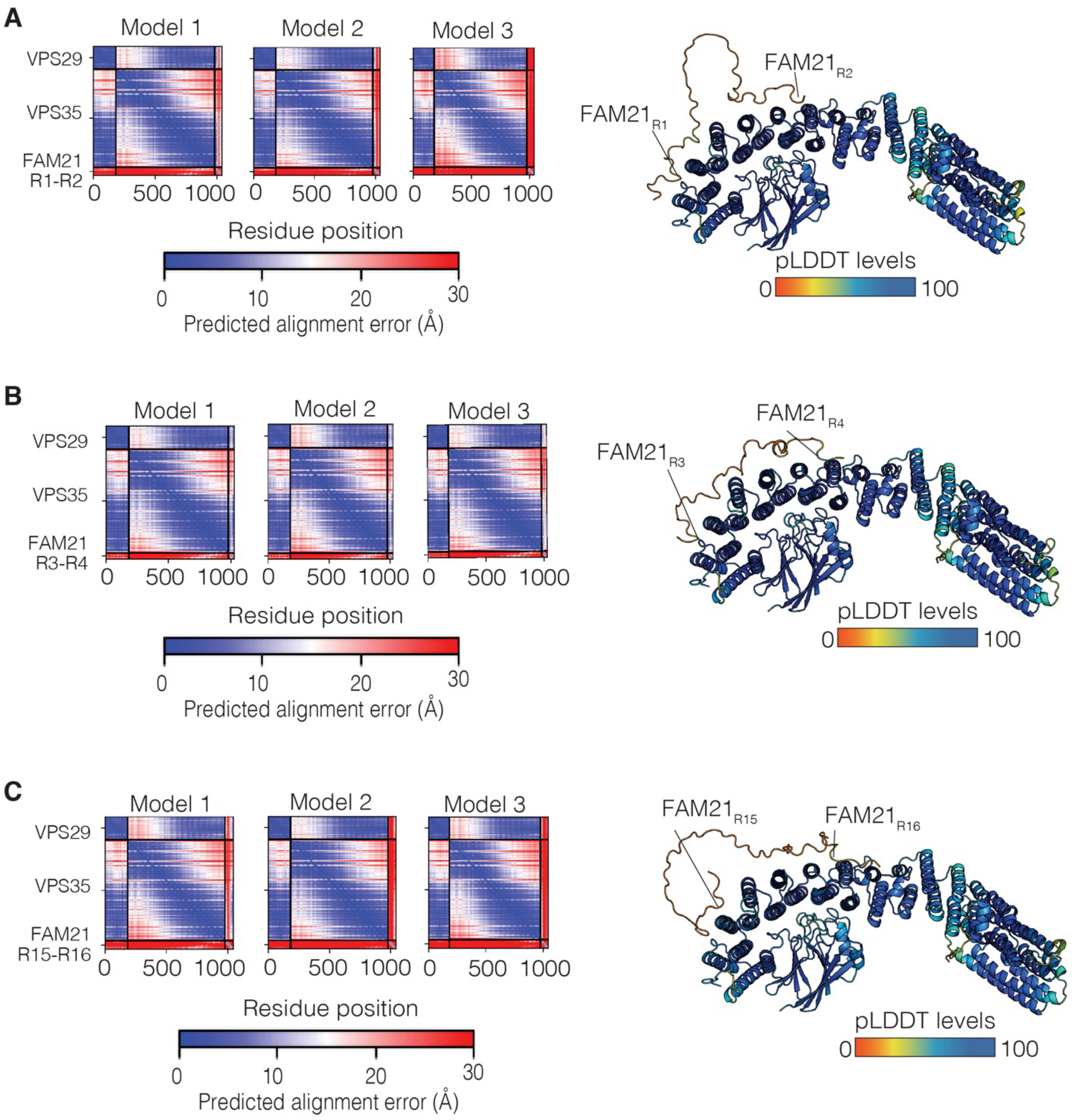


**Figure S3. Alphafold models of VPS35–VPS29 in complex with different FAM21 LFa repeats.**
(A) Predicted alignment Error (PAE) plots for the AlphaFold2-predicted structures of the human (A) VPS35–VPS29–FAM21R1-R2 complex, (B) VPS35–VPS29–FAM21R3-R4 complex, (C) VPS35–VPS29–FAM21R15-R16 complex. On the right the predicted structures are shown coloured according to the per-residue confidence score (pLDDT) between 0 to 100. Regions showing a low pLDDT score (green and orange) are likely to be unstructured.

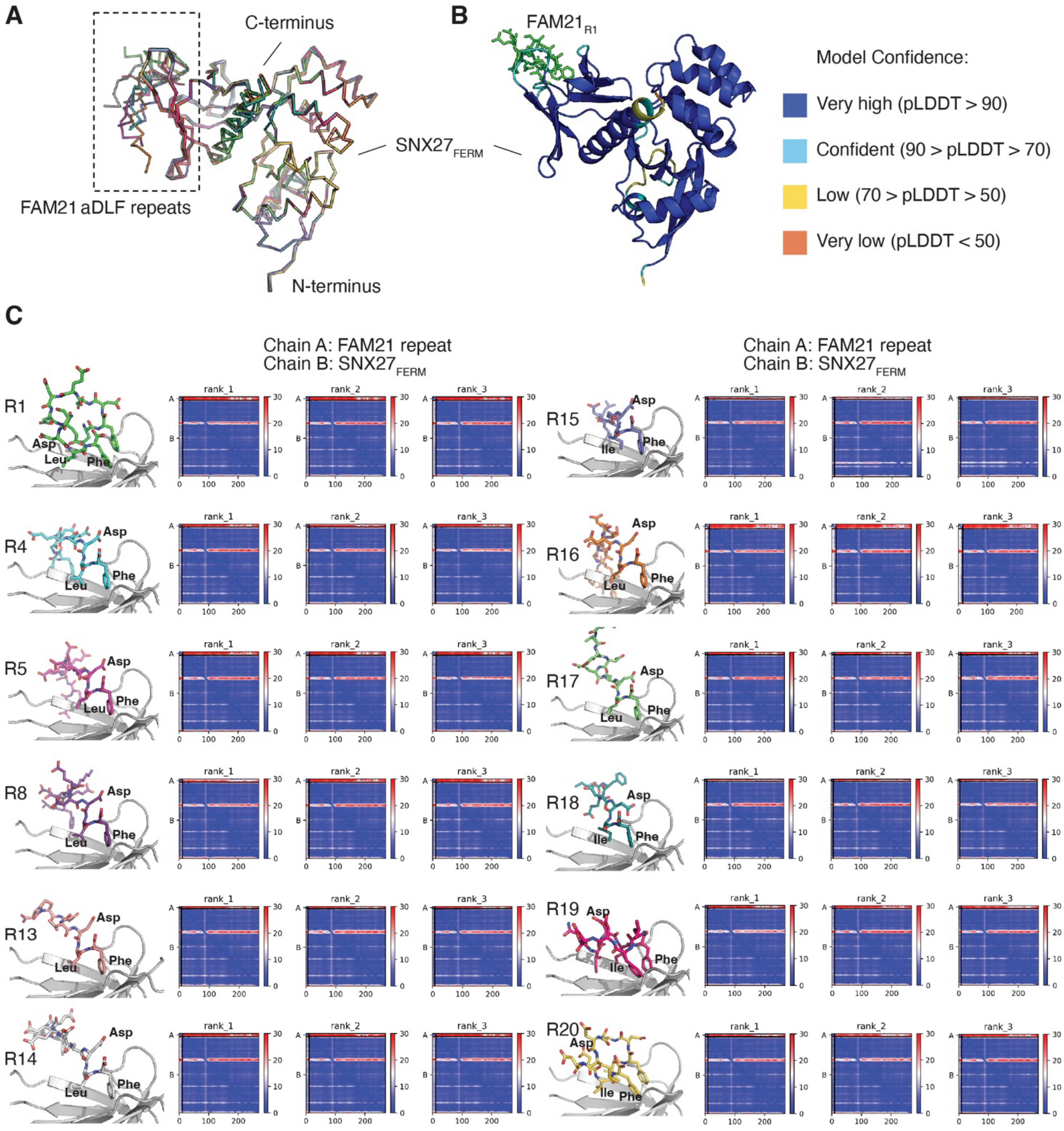


**Figure S4. Alphafold models of SNX27 FERM domain in complex with various FAM21 aDLF sequences.**
(A) Overlay of all AlphaFold2 predicted structures shown as C backbone ribbon. The sequences of the aDLF repeats are shown in Figure 4F. (B) The predicted complex of SNX27FERM bound to FAM21 aDLF sequence from repeat 1. SNX27 is shown in cartoon representation coloured according to pLDDT score. The FAM21R1 peptide is shown in green sticks. (C) Predicted alignment Error (PAE) plots showing the predicted structure of human SNX27FERM–FAM21 aDLF complexes. The top ranked prediction is shown in close up in the left hand panels.

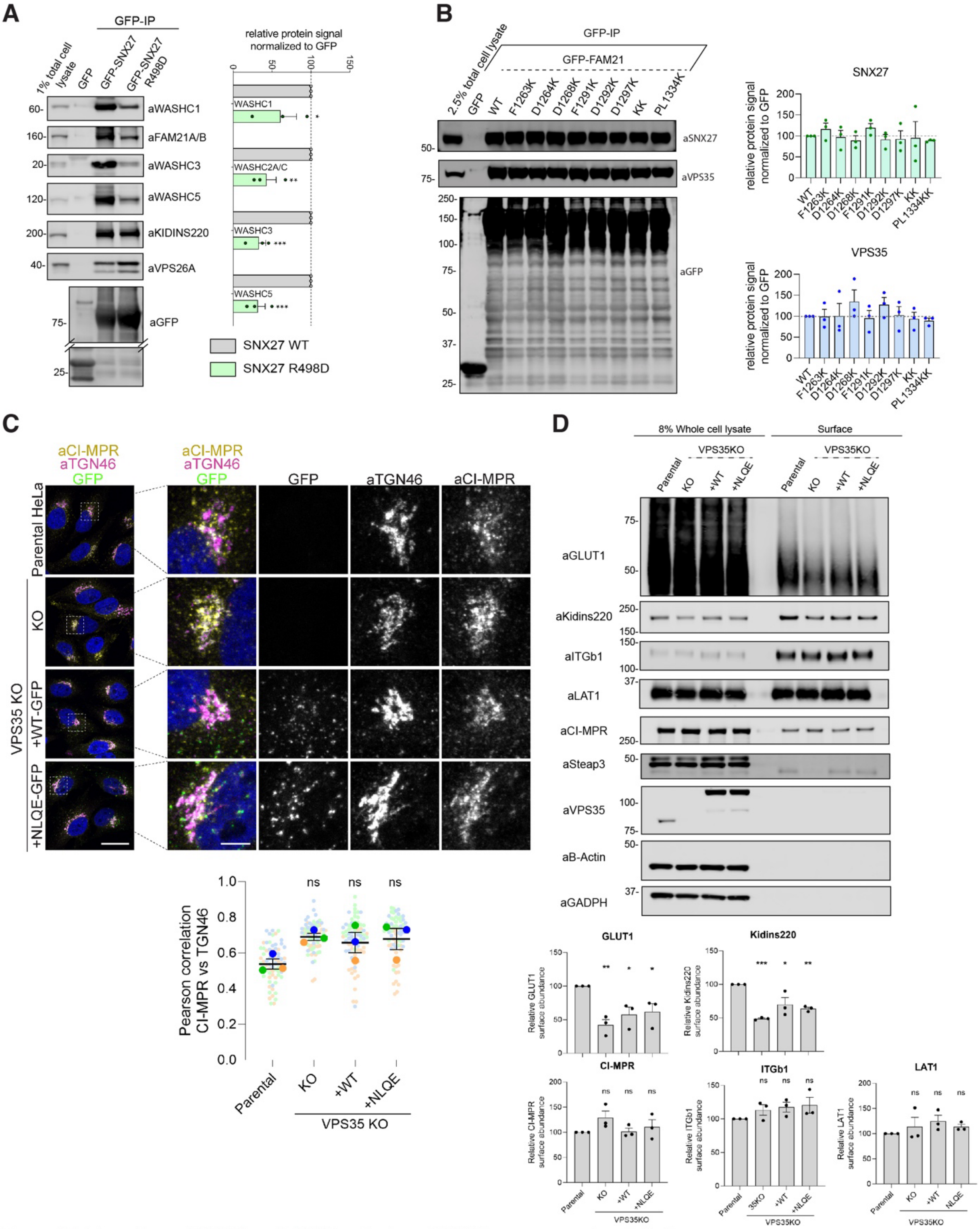

**Figure S5. Interactions of SNX27 and FAM21 mutants and VPS35 rescue experiments in H4 cells.**
(A) Co-IP in HEKs of GFP-SNX27 wild-type and R498D mutant and binding of WASH, Retromer (indicated by subunit VPS26A) and model cargo KIDINS220 assessed by Western blot. (B) Co-IP of GFP-FAM21 and mutants shows that specific mutations in repeats 19-21 do not prevent binding to Retromer or SNX27 in cells. KK is the combination of D1268K and D1297K mutants. (C) CIMPR colocalization with TGN46 is not altered in VPS35 KO, wild-type or NLQE mutant rescued HeLa cells. A total of 70 cells were analyzed in each condition for colocalization between SNX1 and FAM21 across n=3 independent experiments. Scale bars, 25 µm (micrographs) and 5 µm (magnified images). (D) VPS35 KO H4 cells showed reduced cell surface levels of SNX27–Retromer dependent cargos including GLUT1 and KIDINS220. Re-expression and rescue with both GFP-VPS35 wild-type protein and VPS35(NLQE)-GFP is able to partially rescue the cell surface levels of these cargos. Band intensities for all blots were quantified using Odyssey software and normalized to GFP expression. 1-way ANOVA with Dunnett's multiple comparisons test. Error bars represent the mean, s.d. $^{*}P < 0.05$, $^{**}P < 0.01$, $^{***}P < 0.001$, $^{****}P < 0.0001$.

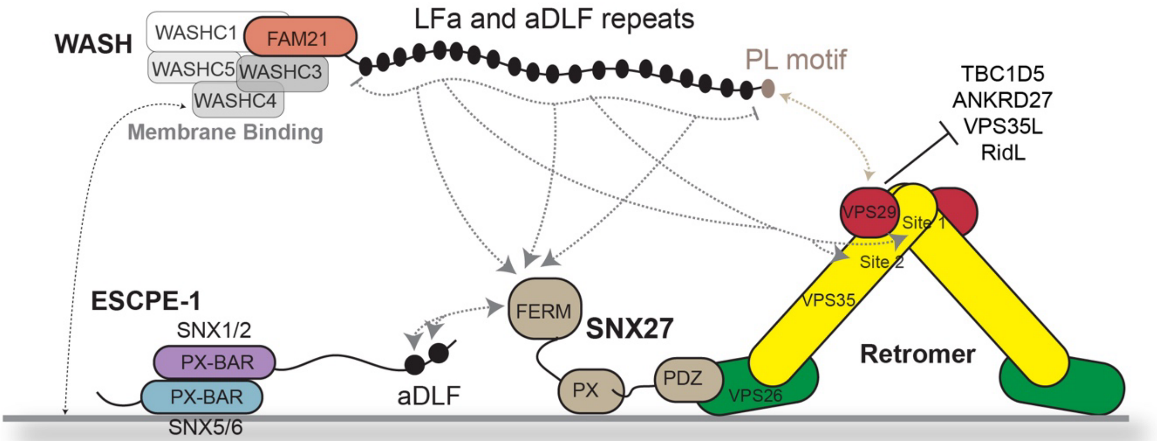


**Figure S6. Summary and proposed model for the cooperative recruitment of FAM21 by Retromer, SNX27 and other endosomal proteins.**

The WASH subunit FAM21 has 21 LFa repeated elements in its extended tail that are rich in acidic, isoleucine and phenylalanine residues. LFa sequences possess two [L/I]F motifs interspersed by an acidic stretch ([L/I]F$[D/E]_{3\text{-}9}$[L/I]F). These bind two conserved sites towards the C-terminus of the VPS35 subunit of Retromer. The last repeated element of FAM21 has a specific PL dipeptide sequence that promotes interaction with Retromer subunit VPS29. This would be mutually exclusive of binding to other VPS29 interactors such as TBC1D5, ANKRD27, VPS35L and bacterial effector RidL. aDLF sequences are a subset of the LFa repeats with an acidic stretch prior to a D[L/I]F motif that associate with a conserved pocket in the SNX27 FERM domain, and are also found in the SNX1 and SNX2 proteins of the ESCPE-1 complex.

### SUPPLEMENTARY MATERIALS:

**Table S1. Thermodynamic parameters for the binding of FAM21 with Retromer by ITC**

| **Syringe** | **Cell** | $K_d$ **(nM)** | $\Delta H$ **(kcal/mol)** | $\Delta G$ **(kcal/mol)** | $-T\Delta S$ **(kcal/mol)** |
|---|---|---|---|---|---|
| | | | **FAM21 peptides against Retromer** | | |
| $FAM21_{R19}$ | Retromer | 63.5 ± 8.7 | -1.2 ± 0.2 | -5.7 ± 0.1 | -4.5 ± 0.3 |
| $FAM21_{R20}$ | Retromer | | N.B. | | |
| $FAM21_{R21}$ | Retromer | 26.8 ± 4.3 | -2.8 ± 0.6 | -6.2 ± 0.1 | -3.4 ± 0.7 |
| | | | **FAM21 peptides against mVps29 – $mVps35_{481-796}$** | | |
| $FAM21_{R19}$ | mVps29 – $mVps35_{481-796}$ | 61.1 ± 6.4 | -1.9 ± 1.4 | -5.8 ± 0.1 | -3.8 ± 1.3 |
| $FAM21_{R20}$ | mVps29 – $mVps35_{481-796}$ | | N.B. | | |
| $FAM21_{R21}$ | mVps29 – $mVps35_{481-796}$ | 31.4 ± 1.1 | -2.6 ± 1.0 | -6.2 ± 0.1 | -3.6 ± 1.2 |
| | | | **FAM21 peptides against mVps29** | | |
| $FAM21_{R19}$ | mVps29 | | N.B. | | |
| $FAM21_{R20}$ | mVps29 | | N.B. | | |
| $FAM21_{R21}$ | mVps29 | 26.6 ± 2.6 | -2.9 ± 0.1 | -6.3 ± 0.1 | -3.4 ± 0.2 |

**Table S2. Summary of crystallographic structure determination statistics of Retromer subcomplex with FAM21 repeats.**

| Data collection statistics | mVPS29 - mVPS35$_{481-796}$ - FAM21$_{R20}$ | mVPS29 - mVPS35$_{481-796}$ - FAM21$_{R21}$ | mVPS29 - FAM21$_{R21}$ |
|---|---|---|---|
| PDB ID | 8TTC | 8TTA | 8TTD |
| Space group | P4$_3$2$_1$2 | C121 | P3$_2$21 |
| Resolution (Å) | 46.90 - 3.01 (3.20 - 3.01) | 49.31 - 3.46 (3.79 - 3.46) | 41.28 - 2.01 (2.06 - 2.01) |
| a, b, c (Å) | 88.76, 88.76, 328.29 | 123.73, 165.58, 68.00 | 43.34, 43.34, 165.14 |
| α, β, γ (°) | 90.00 90.00 90.00 | 90.00, 117.51, 90.00 | 90.00, 90.00, 120.00 |
| Total observations | 348,827 (54,935) | 110,783 (26,173) | 239,643 (17,499) |
| Unique reflections | 27,047 (4,201) | 15,793 (3,693) | 12,765 (911) |
| Completeness (%) | 99.6 (97.9) | 99.3 (97.6) | 99.9 (99.0) |
| $R_{merge}^{+}$ | 0.137 (1.588) | 0.342 (1.233) | 0.052 (0.475) |
| $R_{pim*}$ | 0.040 (0.451) | 0.139 (0.494) | 0.016 (0.152) |
| CC1/2 | 0.998 (0.812) | 0.993 (0.762) | 1.000 (0.972) |
| $<I/\sigma(I)>$ | 11.7 (1.3) | 4.5 (1.3) | 25.4 (4.5) |
| Multiplicity | 12.9 (13.1) | 7.0 (7.1) | 18.8 (19.2) |
| Molecule/asym | 2 | 2 | 1 |
| **Refinement statistics** | | | |
| $R_{work}/R_{free}$ (%) ¶# | 24.90/27.40 | 22.22/25.39 | 18.81/24.41 |
| No. protein atoms | 7747 | 7908 | 1507 |
| Waters | 17 | 10 | 26 |
| Wilson B (Å$^2$) | 90.59 | 77.56 | 42.24 |
| Average B (Å$^2$)^ | 101.87 | 83.95 | 59.45 |
| Protein | 102.13 | 84.00 | 59.53 |
| ligands | 63.10 | 85.78 | 96.35 |
| Water | 83.53 | 41.92 | 50.70 |
| rmsd bonds (Å) | 0.006 | 0.006 | 0.007 |
| rmsd angles (°) | 1.11 | 1.08 | 1.08 |
| Ramachandran plot: Favored/outliers (%) | 93.28/1.37 | 93.42/0.93 | 94.02/0.54 |

Values in parentheses refer to the highest resolution shell. $^{+}R_{merge} = \Sigma|I - <I>| / \Sigma<I>$, where $I$ is the intensity of each individual reflection. *$R_{pim}$ indicates all $I^{+}$ & $I^{-}$. ¶$R_{work} = \Sigma h|F_o - F_c| / \Sigma_l| F_o|$, where $F_o$ *and* $F_c$ are the observed and calculated structure-factor amplitudes for each reflection $h$. #$R_{free}$ was calculated with 10% of the diffraction data selected randomly and excluded from refinement. ^Calculated using Baverage.

**Table S3. Thermodynamic parameters for the binding of FAM21 with SNX27 by ITC**

| Syringe | Cell | $K_d$ (μM) | $\Delta H$ (kcal/mol) | $\Delta G$ (kcal/mol) | $-T\Delta S$ (kcal/mol) |
|---|---|---|---|---|---|
| | | **$FAM21_{R19-21}$ against SNX27 proteins** | | | |
| $FAM21_{R19-21}$ | $hSNX27_{FL}$ | 5.70 ± 1.94 | -1.51 ± 0.45 | -7.18 ± 0.21 | -5.67 ± 0.25 |
| $FAM21_{R19-21}$ | $hSNX27_{FERM}$ | 4.97 ± 1.32 | -2.50 ± 0.25 | -7.25 ± 0.16 | -4.75 ± 0.41 |
| | | **FAM21 peptides against $hSNX27_{FL}$** | | | |
| $FAM21_{R5}$ | $hSNX27_{FL}$ | 23.55 ± 2.19 | -1.28 ± 0.15 | -6.32 ± 0.05 | -5.04 ± 0.10 |
| $FAM21_{R8}$ | $hSNX27_{FL}$ | 68.80 ± 1.27 | -2.12 ± 0.04 | -5.69 ± 0.01 | -3.57 ± 0.05 |
| $FAM21_{R17}$ | $hSNX27_{FL}$ | 70.55 ± 0.64 | -2.10 ± 0.57 | -5.67 ± 0.01 | -3.57 ± 0.58 |
| $FAM21_{R19}$ | $hSNX27_{FL}$ | 6.96 ± 0.12 | -2.22 ± 0.17 | -7.04 ± 0.01 | -4.82 ± 0.18 |
| $FAM21_{R20}$ | $hSNX27_{FL}$ | 19.00 ± 0.28 | -1.70 ± 0.28 | -6.45 ± 0.01 | -4.75 ± 0.29 |
| $FAM21_{R20}$ mutant | $hSNX27_{FL}$ | | No binding detected | | |
| $FAM21_{R21}$ | $hSNX27_{FL}$ | | No binding detected | | |
| | | **FAM21 peptides against $hSNX27_{FERM}$** | | | |
| $FAM21_{R5}$ | $hSNX27_{FERM}$ | 29.20 ± 0.28 | -3.05 ± 0.62 | -6.19 ± 0.01 | -3.14 ± 0.61 |
| $FAM21_{R8}$ | $hSNX27_{FERM}$ | 42.50 ± 2.97 | -2.87 ± 0.19 | -5.97 ± 0.04 | -3.11 ± 0.23 |
| $FAM21_{R17}$ | $hSNX27_{FERM}$ | 31.70 ± 0.57 | -3.11 ± 0.92 | -6.14 ± 0.01 | -3.03 ± 0.93 |
| $FAM21_{R19}$ | $hSNX27_{FERM}$ | 38.35 ± 1.34 | -3.33 ± 0.52 | -6.03 ± 0.02 | -2.70 ± 0.49 |
| $FAM21_{R20}$ | $hSNX27_{FERM}$ | 25.55 ± 0.35 | -3.00 ± 0.29 | -6.27 ± 0.01 | -3.27 ± 0.28 |
| $FAM21_{R21}$ | $hSNX27_{FERM}$ | | No binding detected | | |
| | | **$FAM21_{R19}$ peptide against $hSNX27_{FL}$ mutants** | | | |
| $FAM21_{R19}$ | $hSNX27_{FL}$ R437D | | No binding detected | | |
| $FAM21_{R19}$ | $hSNX27_{FL}$ K495D | 58.30 ± 1.41 | -2.33 ± 0.34 | -5.78 ± 0.01 | -3.45 ± 0.35 |
| $FAM21_{R19}$ | $hSNX27_{FL}$ K496D | 33.53 ± 1.48 | -2.97 ± 1.53 | -4.99 ± 1.96 | -3.14 ± 1.52 |
| $FAM21_{R19}$ | $hSNX27_{FL}$ R498D | | No binding detected | | |
| $FAM21_{R19}$ | $hSNX27_{FL}$ K501D | 38.80 ± 0.28 | -2.14 ± 0.95 | -6.02 ± 0.00 | -3.88 ± 0.95 |

**Table S4. Summary of crystallographic structure determination statistics for $SNX27_{FERM}$ and FAM21 repeats.**

| **Data collection statistics** | **$hSNX27_{FERM}$ - $FAM21_{R15}$** | **$hSNX27_{FERM}$ - $FAM21_{R19}$** | **$hSNX27_{FERM}$ - $FAM21_{R20}$** |
|---|---|---|---|
| PDB ID | 8TTT | 8TTU | 8TTV |
| Space group | $P2_12_12_1$ | $P2_12_12_1$ | $P2_12_12_1$ |
| Resolution (Å) | 42.95– 2.33 | 37.54 – 2.36 | 42.91 – 2.00 |
| | (2.41 – 2.33) | (2.45 – 2.36) | (2.05 – 2.00) |
| a, b, c (Å) | 37.57, 74.37, 105.23 90.00, | 37.54, 74.50, 102.53 | 37.97, 74.20, 105.19 |
| α, β, γ (°) | 90.00, 90.00 | 90.0, 90.0, 90.0 | 90.0, 90.0, 90.0 |
| Total observations | 88510 (5906) | 64599 (6238) | 139518 (8637) |
| Unique reflections | 12909 (940) | 12319 (1195) | 20806 (1385) |
| Completeness (%) | 97.6 (75.9) | 98.8 (93.5) | 99.5 (94.2) |
| $R_{merge}^{+}$ | 0.203 (1.524) | 0.097 (0.545) | 0.240 (2.161) |
| $R_{pim*}$ | 0.083 (0.632) | 0.048 (0.263) | 0.101 (0.943) |
| CC1/2 | 0.995 (0.379) | 0.995 (0.801) | 0.989 (0.205) |
| $\langle I/\sigma(I)\rangle$ | 7.9 (1.1) | 8.4 (1.7) | 6.7 (1.0) |
| Multiplicity | 6.9 (6.3) | 5.2 (5.2) | 6.7 (6.2) |
| Molecule/asymmetry | 1 | 1 | 1 |
| **Refinement statistics** | | | |
| $R_{work}/R_{free}$ (%) ¶# | 19.05/25.04 | 20.04/25.26 | 20.64/26.38 |
| No. protein atoms | 2187 | 2192 | 2184 |
| Waters | 45 | 28 | 116 |
| Wilson B (Å$^2$) | 39.05 | 47.82 | 27.32 |
| Average B (Å$^2$)^ | 44.23 | 55.02 | 36.53 |
| Protein | 44.22 | 54.95 | 36.51 |
| Ligands | 49.37 | 64.36 | 39.68 |
| Water | 41.71 | 51.53 | 36.70 |
| rmsd bonds (Å) | 0.009 | 0.009 | 0.009 |
| rmsd angles (°) | 1.03 | 0.98 | 0.98 |
| Ramachandran plot: | | | |
| Favored/outliers (%) | 95.63/0.79 | 94.86/1.58 | 94.80/2.40 |

Values in parentheses refer to the highest resolution shell. $^{+}R_{merge} = \Sigma|I - \langle I\rangle| / \Sigma\langle I\rangle$, where $I$ is the intensity of each individual reflection. *$R_{pim}$ indicates all $I^+$ & $I^-$. ¶$R_{work} = \Sigma h|F_o - F_c| / \Sigma_l| F_o|$, where $F_o$ *and* $F_c$ are the observed and calculated structure-factor amplitudes for each reflection $h$. #$R_{free}$ was calculated with 10% of the diffraction data selected randomly and excluded from refinement. ^Calculated using Baverage.

**Table S5. Key Resources**

| REAGENT or RESOURCE | SOURCE or REFERENCE | IDENTIFIER |
|---|---|---|
| **Bacterial Strains** | | |
| *E. coli* BL21 (DE3) | Invitrogen | Catalogue: C600003 |
| **Chemicals** | | |
| Benzamidine hydrochloride hydrate | Sigma Aldrich | B6506 |
| Deoxyribonuclease I (DNase I) | Sigma Aldrich | DN25 |
| Ampicillin, Sodium Salt | Astral Scientific | A-1414-25g |
| Kanamycin Sulfate | Astral Scientific | K-1022-25g |
| Talon® resin | Clontech | 635503 |
| Glutathione Sepharose 4B | GE Healthcare | GEHE17-0756-0 |
| Glutathione (reduced form) | Novachem | Catalogue: 077-02016 |
| Isopropyl β-D-1-thiogalactopyranoside | Bioline | Catalogue: BIO-37036 |
| 2-Mercaptoethanol (β-ME) | Sigma Aldrich | Catalogue: M6250-100ML |
| Triton-X100 | Sigma Aldrich | X100-500ML |
| cOmpleteTM EDTA-free protease inhibitor tablet | Sigma Aldrich | Catalogue: 4693159001 |
| EZ-Link Sulfo-NHS-SS Biotin | Thermo Fisher Scientific | Catalogue: 21217 |
| **Synthesised peptides** | | |
| FAM21-R5$_{536-552}$ GLFSDEEDSEDLFSSQS | Genscript | N/A |
| FAM21-R8$_{662-678}$ TSLFEEDEEDDLFAIAK | Genscript | N/A |
| FAM21-R15$_{1124-1140}$ RGEADLFDSGDIFSTGT | Genscript | N/A |
| FAM21-R17$_{1197-1213}$ NPFPLLEDEDDLFTDQK | Genscript | N/A |
| FAM21-R19$_{1261-1274}$ NLFDDNIDIFADLT | Genscript | N/A |
| FAM21-R20$_{1289-1302}$ SIFDDDMDDIFSSG | Genscript | N/A |
| FAM21-R20$_{1289-1302\ (IF/SS)}$ S**SS**DDDMDD**SS**SSG | Genscript | N/A |
| FAM21-R21$_{1328-1341}$ SNIFDDPLNAFGGQ | Genscript | N/A |
| **Recombinant DNA** | | |
| Plasmid: pET-28a human SNX27$_{FL}$ | Gene Universal | N/A |
| Plasmid: pET-28a human SNX27$_{FERM}$ | Gene Universal | N/A |
| Plasmid: pGEX4T-2 human FAM21$_{R19-21}$ | Gene Universal | |
| Plasmid: pET-28a human SNX27$_{FL}$ (R437D) | Gene Universal | N/A |
| Plasmid: pET-28a human SNX27$_{FL}$ (K495D) | Gene Universal | N/A |
| Plasmid: pET-28a human SNX27$_{FL}$ (K496D) | Gene Universal | N/A |
| Plasmid: pET-28a human SNX27$_{FL}$ (R498D) | Gene Universal | N/A |
| Plasmid: pET-28a human SNX27$_{FL}$ (K501D) | Gene Universal | N/A |
| Plasmid: pEGFP-c1-SNX27 WT and mutants | (2) | N/A |
| Plasmid: pEGFP-n1-VPS35 WT and mutants | This study | N/A |
| Plasmid: pEGFP-c1-Fam21 Tail WT, deletions and mutants | This study | N/A |
| Plasmid: pXLG3-VPS35-GFP WT and K555N, K556L, K559Q | This study | N/A |
| **Software and Algorithms** | | |
| Pymol (version 2.5.4) | Schrodinger, USA. | https://pymol.org/2/ |

| | | |
|---|---|---|
| Chimera (version 1.4.0) | Resource for Biocomputing, Visualization, and Informatics, University of California, USA | https://www.cgl.ucsf.edu/chimera/ |
| Prism 7 | GraphPad | https://www.graphpad.com/features |
| **Other** | | |
| HiLoad® 16/600 Superdex 200 PG | GE Healthcare | Catalogue: 28-9893-35 |
| Mono Q® 10/100 GL | GE Healthcare | Catalogue: 17-5167-01 |
| Amicon™ ultrafiltration devices | Merck | Catalogue: 900000042327 (10K)<br>Catalogue: 900000008078 (30K) |
| SDS-PAGE BOLT gels | Thermo Fisher Scientific | Catalogue: NW04122BOX (12-well) |
| NuPAGE 4-12% gels | Thermo Fisher Scientific | Catalogue: NP0336 |
| NuPAGE MOPS SDS running buffer | Thermo Fisher Scientific | Catalogue: NP000102 |
| ChromoTek GFP-Trap Agarose beads | ProteinTech | Catalogue: gta |
| Streptavidin Sepharose beads | Cytiva | Catalogue: 90100484 |
